# NeuroPilot: Longitudinal intravital neural imaging across scales and modalities through an agentic platform

**DOI:** 10.64898/2026.10.18.759043

**Authors:** Guoxun Zhang, Yuhong Luo, Furong Ju, Dengyu Liu, Dong Zhang, Changliang Guo, Jiaying Han, Shan Qiu, Tianyun Wang, Jun Wang, Mingyang Ma, Bo Li, Yanjun Jiang, Pengfei Wei, Xiaodong Liu, Nan Chi, Ziwei Li

**Affiliations:** College of Future Information Technology, Fudan University, Shanghai 200433, China; Department of Anaesthesia and Intensive Care, The Chinese University of Hong Kong, Hong Kong SAR, China; Department of Pharmaceutical Chemistry, University of California, San Francisco, CA 94143; Department of Automation, Tsinghua University, Beijing 100084, China; PKU-Nanjing Institute of Translational Medicine, Nanjing Raygen Health, Nanjing 211899, China; Academy of Advanced Interdisciplinary Study, Peking University, Beijing 100871, China; State Key Laboratory of Membrane Biology, Beijing Key Laboratory of Cardiometabolic Molecular Medicine, Institute of Molecular Medicine, School of Future Technology, Center for Life Sciences, Peking University, Beijing 100871, China; Department of Neurology, University of California, San Francisco, CA 94143; School of Psychology, Shanghai Jiao Tong University, Shanghai 200030, China; School of Biological Science and Medical Engineering, State Key Laboratory of Digital Medicine, Southeast University, Nanjing 211189, China

## Abstract

Longitudinal intravital imaging is essential for understanding how neural populations evolve across behavioral states, yet converting large-scale recordings into biological insight remains a fundamental challenge because neural signals are degraded by noise, non-rigid motion, and heterogeneous recording conditions. Here we introduce NeuroPilot, an agentic framework that transforms neural imaging analysis into a closed-loop, context-aware process, integrating a multimodal large language model with callable computational imaging modules to adaptively coordinate workflows from raw recordings to population-level readouts. Across heterogeneous imaging and experimental regimes, NeuroPilot restores motion-degraded recordings and supports consistent neural signal recovery, detection, and tracking in challenging deep-brain, freely moving, and longitudinal settings. These capabilities enable neuronal population tracking across spinal stimulation paradigms, deep-brain imaging, and continuous 24-hour mPFC recordings aligned with spontaneous sleep-wake behavior. By converting heterogeneous intravital recordings into stable population activity maps, NeuroPilot supports behavioral decoding and reveals structured neural dynamics across behavioral states and transitions. NeuroPilot establishes a scalable agentic paradigm for neural imaging analysis, expanding experimental access to the measurement, visualization, and interpretation of neural population activity.

## Introduction

Deciphering how neural ensembles encode behavior and internal states across time remains a central challenge in neuroscience^1,2^. Brain activity is inherently dynamic and distributed, unfolding across three-dimensional circuits as animals engage in ongoing behavior, transition between internal states, and undergo physiological or pathological change^3–5^. Capturing these dynamics requires following the same neuronal populations across spatial and temporal scales: from local cellular interactions to circuit-wide activity patterns, and from transient state changes to long-term remodeling^6,7^. Intravital calcium imaging has brought this goal increasingly within reach, extending cellular-resolution recording across anatomical regions, imaging depths, behavioral contexts, and prolonged observation windows in living animals^8–13^. Despite these advances, as neural imaging moves toward larger, deeper, longer, and more naturalistic recordings, analysis has emerged as a central constraint: extracting stable neuronal footprints, faithful calcium traces, and interpretable population dynamics from raw, heterogeneous recordings remains a major bottleneck^14,15^.

This bottleneck arises because the recording regimes essential for naturalistic and longitudinal neural imaging often yield data that are difficult to convert into reliable neural activity readouts. Long-duration recordings are constrained by trade-offs among photon budget, shot-noise limits, and phototoxicity, often resulting in weak signals, photobleaching, baseline drift, and intermittent signal loss^16–18^. Recordings in freely moving animals, under sensory stimulation, in deep tissue, or using miniaturized microscopes further introduce non-rigid motion, reduced photon-collection efficiency, and mixed spatial signals^19–22^. These difficulties are amplified when experiments aim to construct neural data matrices across stimuli, behavioral states, sessions, and disease progression. In such settings, longitudinal analysis requires integrating recordings across conditions and days into a coherent record while preserving the identity of individual neurons across sessions^7,23,24^.

Reliable neural imaging analysis depends not only on the robustness of individual algorithms, but also on how these algorithms are coordinated and adapted across changing recording conditions. Existing tools address key components of neural imaging analysis, including denoising, motion correction, segmentation, trace extraction, dimensionality reduction, and behavioral decoding^25–28^. However, individual methods often lose robustness when confronted with severe motion, weak or unstable signals, heterogeneous imaging modalities, or long-term changes in tissue and optical conditions. Moreover, these tools are typically assembled into manually configured, dataset-specific workflows that require extensive intervention, impose substantial processing burdens, and adapt poorly when data characteristics change. This challenge is compounded by the strong interdependence among processing stages, which allows uncertainties to propagate throughout the analytical workflow. Analysis also remains largely separated from image acquisition, preventing processing decisions from responding directly to data quality and experimental context. Addressing these limitations therefore requires an integrated and adaptive framework that combines robust computational operators with intelligent workflow coordination, links restoration, motion correction, cell identification and tracking, and downstream interpretation, and preserves a traceable path from raw recordings to population-level neural readouts.

Here, we introduce NeuroPilot, an agentic framework for multimodal intravital neural imaging analysis orchestrated by the large language model (LLM). NeuroPilot treats analysis as an iterative, closed-loop process rather than a fixed linear pipeline, coordinating denoising, dense non-rigid motion correction, cell segmentation, trace extraction, quality assessment, and population-level analysis. To support stabilization under complex imaging conditions, we developed PyLoReg-Net, a pixel-wise non-rigid registration network that estimates dense displacement fields to compensate for spatially heterogeneous motion and deformation. The framework is available as a cloud-accessible platform through an interactive web interface (https://neuropilot.it.com/). We first validate NeuroPilot on paired low-and high-SNR two-photon recordings. We then test its robustness across one-, two-, and three-photon calcium imaging, deep-brain and multiplexed recordings, and longitudinal experiments spanning repeated stimulation paradigms, cross-day tracking of spinal cord neurons, and continuous 24-hour mPFC imaging (Figure 1A)^9,29–31^. In spinal cord imaging, NeuroPilot tracks matched neurons across freely moving behavior, sensory stimulation, and repeated post-injury sessions, linking single-cell activity across conditions and days. In continuous 24-hour mPFC imaging, NeuroPilot maintains stable population readouts throughout spontaneous sleep-wake behavior, supporting behavioral decoding and latent-dynamics analysis of state transitions. Together, these results establish NeuroPilot as an adaptive, traceable platform for converting raw intravital recordings into structured representations of neural activity across spatial and temporal scales (Figure 1B).

**Figure 1.**
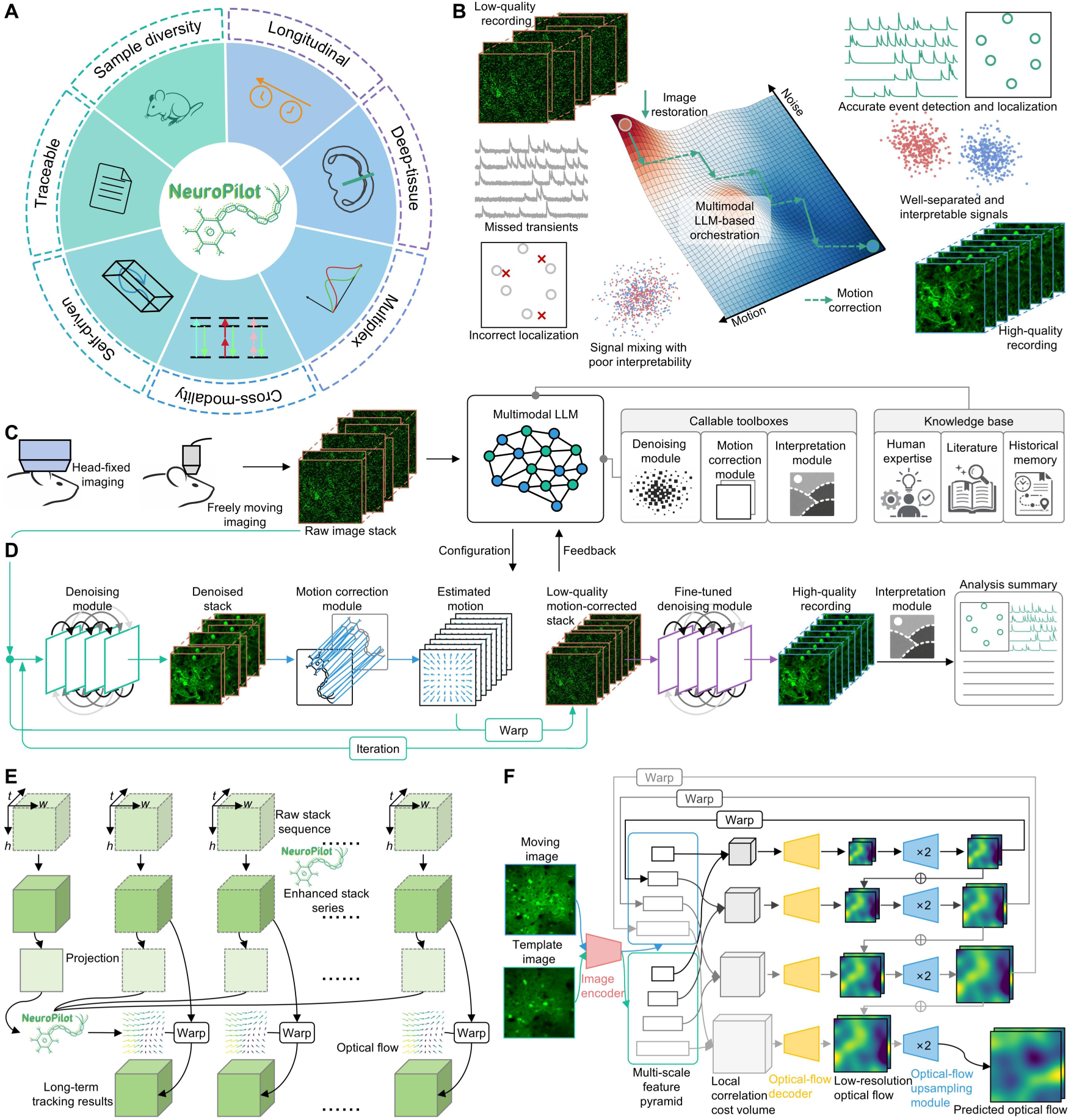
NeuroPilot establishes a closed-loop agentic framework for multimodal intravital neural imaging. (A) Overview of NeuroPilot capabilities. NeuroPilot provides automated and traceable workflows across sample diversity, longitudinal tracking, deep-tissue imaging, multiplexing, and one-to three-photon modalities. (B) Problem setting and motivation for NeuroPilot. NeuroPilot leverages multimodal LLM orchestration to recover neural signals from noisy and motion-corrupted recordings, supporting reliable activity readout across heterogeneous imaging conditions. (C) Architecture of NeuroPilot. Heterogeneous neural recordings are routed to a multimodal LLM, which configures processing workflows by coordinating callable modules for image restoration, registration, and interpretation. A knowledge base integrating human expertise, literature-derived priors, and historical processing records supports context-aware module selection and iterative refinement. (D) Autonomous iterative processing pipeline. Intermediate outputs are evaluated and fed back to the LLM-orchestrated workflow, which revises processing strategies until predefined quality criteria are met. This closed-loop procedure progressively improves spatial stability, structural visibility, and signal fidelity, producing analysis-ready recordings for downstream neuronal activity extraction. (E) Hierarchical cross-session processing. Recordings acquired across sessions or conditions are enhanced and projected to estimate optical flow. The resulting deformation fields map each recording into a shared reference frame, enabling cross-session correspondence and longitudinal tracking of neuronal populations. (F) Coarse-to-fine registration in PyLoReg-Net. Moving and template images are encoded into multi-scale feature pyramids. At each scale, local correlation cost volumes estimate residual optical flow, which is integrated with upsampled estimates from coarser levels to generate the final dense deformation field.

## Results

### NeuroPilot establishes a closed-loop agentic framework for scalable neural imaging analysis

At the system level, NeuroPilot ingests raw image stacks acquired using head-fixed microscopes or freely moving miniscopes and uses a multimodal LLM as the planner to configure callable toolboxes for self-supervised denoising, non-rigid motion correction, and neural activity interpretation (Figure 1C). The planner is coupled to a knowledge base that incorporates human expertise, literature-derived priors, and historical processing records. This design allows processing choices to be contextualized by the properties of each recording, including imaging parameters, recording conditions, behavioral paradigms, and experimental design. Instead of applying the same fixed sequence of operations to every dataset, NeuroPilot uses intermediate outputs, including restored stacks, deformation fields and quality-control metrics to guide subsequent processing decisions. The resulting workflow provides a traceable path from raw recordings to restored image stacks, neuronal footprints, extracted calcium traces, and summary reports (Video S1).

The computational core of NeuroPilot is a closed-loop restoration strategy that couples self-supervised denoising with non-rigid motion correction (Figure 1D; Methods). In each cycle, the self-supervised denoising module recovers underlying neuronal structures and enhances their spatial features, thereby improving the accuracy of subsequent motion estimation. The motion-correction module then estimates dense deformation fields to register the noisy input stack to a common spatial reference. The corrected stack is fed back into the next round of restoration, allowing signal recovery and spatial alignment to be refined jointly. Final signal restoration is executed only once to prevent over-smoothing caused by repeated denoising. This iterative formulation is designed for recordings in which noise and motion are interdependent: poor signal quality impairs registration, whereas residual motion degrades trace extraction and downstream interpretation. After image restoration, the same agentic workflow dispatches modules for segmentation, activity extraction, quality assessment, and report generation.

To extend NeuroPilot from individual recordings to multi-session datasets, we implemented a hierarchical cross-stack workflow that separates within-session restoration from cross-session alignment (Figure 1E; Methods). Each recording is first restored independently to preserve local signal structure within that stack. Projection images are then generated as registration targets for estimating optical-flow deformation fields across sessions or experimental conditions. Warping each stack into a shared reference frame allows neuronal footprints and calcium traces to be linked across recordings. This design supports cross-condition analysis within a single day, longitudinal neuronal tracking across sessions, and block-wise processing of day-scale continuous recordings.

### PyLoReg-Net estimates dense deformation fields for non-rigid neural image registration

To couple image restoration with motion correction, we developed PyLoReg-Net (pyramid local registration network), a pixel-wise optical-flow registration module for non-rigid motion correction in neural imaging data (Methods). In the NeuroPilot workflow, PyLoReg-Net takes a moving image and a reference image as input and estimates a dense deformation field that warps the recording into a common spatial frame (Figure 1F). The network follows a coarse-to-fine registration strategy: multi-scale feature pyramids are extracted from the moving and reference images, local correlation cost volumes are computed at each pyramid level to establish spatial correspondences, and residual optical-flow fields are progressively refined from coarse to fine resolutions (Figure S1A and Video S1). This design allows global rigid drift and local tissue deformation to be handled within a single dense registration framework.

We trained PyLoReg-Net using a supervised deformation-learning strategy designed for calcium imaging data (Figure S1B-D; Methods). The training corpus included both experimentally acquired neuronal datasets and synthetic calcium imaging datasets, encompassing diverse brain regions, imaging configurations, acquisition conditions, and neuronal morphologies (Figure S1D). To generate paired training examples with known ground-truth deformation fields, we applied simulated affine, elastic, B-spline-like, and Gaussian-mixture-based nonlinear deformations to reference images, followed by noise and blur augmentation (Figure S1B). Each training pair consisted of a deformed moving image and its corresponding undeformed reference image as network inputs, together with the applied deformation field as ground-truth optical-flow supervision (Methods).

After training, PyLoReg-Net was incorporated into the NeuroPilot closed-loop restoration framework. In each iteration, the network estimates dense deformation fields between the moving frames and a reference image, which are used to spatially align the recording. The motion-corrected images then provide a more stable input for subsequent denoising, and the refined images in turn support more accurate motion estimation. This registration step links local image restoration to downstream segmentation, trace extraction, and cross-session alignment. The resulting outputs are compiled into a traceable NeuroPilot report that summarizes raw data properties, restored image quality, motion-correction metrics, neuronal footprints, calcium traces, activity heatmaps, and correlation maps for downstream inspection and analysis (Figure S1E).

### NeuroPilot restores neural structure and calcium dynamics in paired low-SNR recordings

To quantitatively evaluate NeuroPilot restoration, we developed a dedicated dual-channel two-photon microscopy system that acquired paired low- and high-SNR recordings from the same field of view (FOV), with the high-SNR channel serving as an experimentally matched reference (Methods). Fluorescence emission was split into low- and high-signal detection paths using a beamsplitter, which was calibrated to yield an approximately 1:10 signal ratio between the paired image streams (Figure S2A). This configuration generated synchronized recordings with distinct signal levels but matched biological dynamics. The high-SNR recordings were motion corrected to generate reference data^32^, whereas the corresponding low-SNR recordings were processed by NeuroPilot (Figure S2B). This design provided a controlled benchmark for assessing whether NeuroPilot restoration preserved both image structure and calcium activity dynamics.

We first applied this paired benchmark to an FOV containing neuronal somata and dendritic processes. In the raw low-SNR recordings, neuronal somata and neurites were partially obscured by background noise, while temporal kymographs showed disrupted spatial continuity attributable to photon noise and tissue motion. NeuroPilot processing recovered previously obscured somatic and neuritic features and restored spatial continuity in the kymographs, yielding structural patterns that were more consistent with the paired high-SNR reference recordings (Figures 2A-2C). Calcium traces extracted from NeuroPilot-restored recordings better matched the high-SNR reference traces than traces extracted from the raw low-SNR data, including the timing of transient events highlighted in representative neurons (Figures 2D and 2E; Methods). Across 115 segmented neurons, NeuroPilot increased the Pearson correlation between extracted calcium traces and the paired high-SNR reference from 0.382 ± 0.011 to 0.774 ± 0.010 (mean ± SEM; two-sided paired t-test, *P* < 0.0001; Figure 2F; Methods) and increased trace SNR from 1.89 ± 0.03 dB to 6.39 ± 0.13 dB (mean ± SEM; two-sided paired t-test, *P* < 0.0001; Figure 2G). Estimated displacement fields further showed spatially heterogeneous motion in the low-SNR recordings, with local displacement magnitudes reaching 25 μm, supporting the need for non-rigid, motion-aware restoration (Figure 2H).

**Figure 2.**
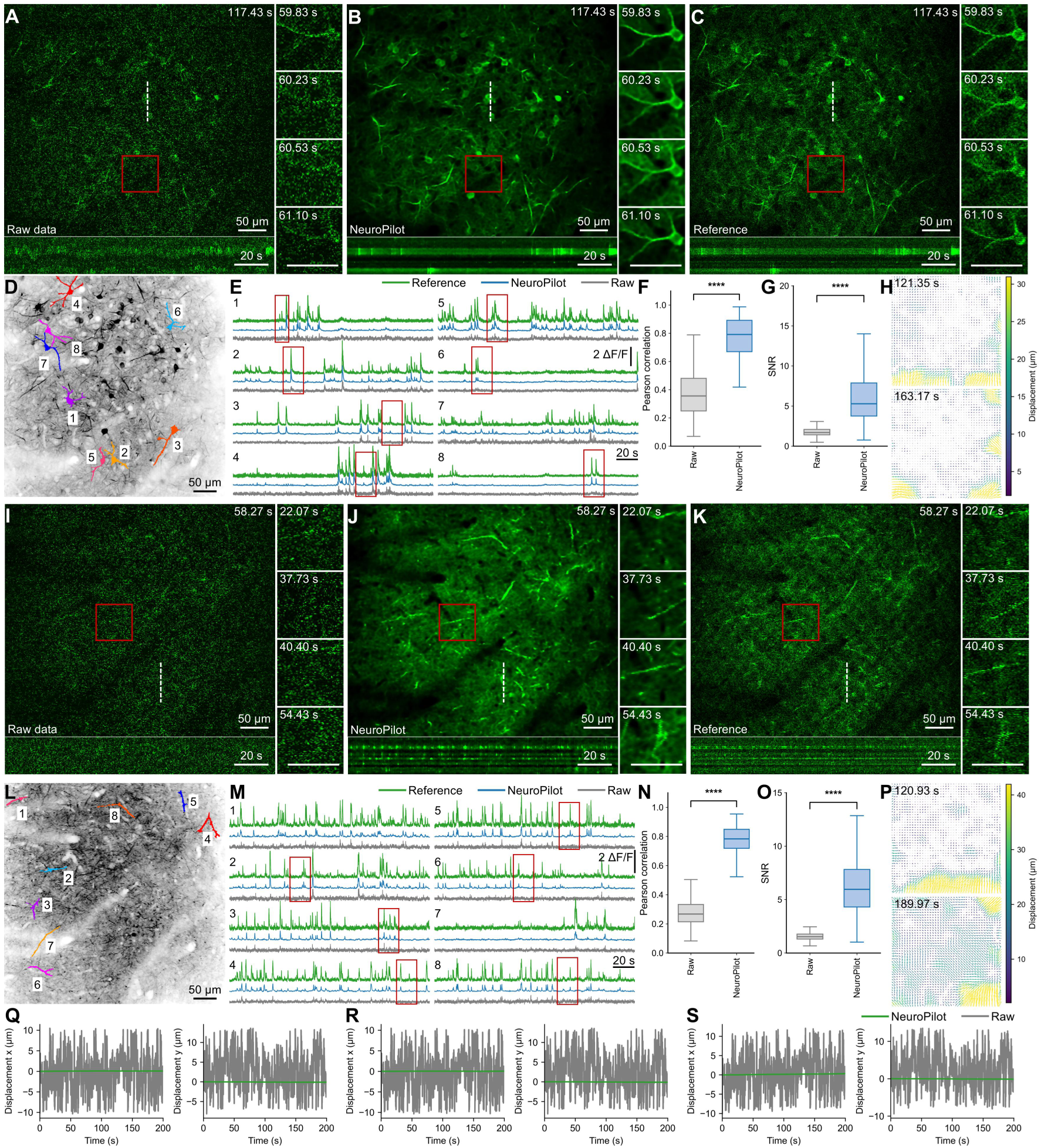
NeuroPilot restores calcium activity from recordings jointly degraded by noise and motion. (A-C) Example FOV containing neuronal somata and dendritic processes. Representative images show the raw low-SNR recording (A), the corresponding NeuroPilot-restored result (B), and the paired high-SNR reference (C). Red boxes indicate regions enlarged in the corresponding insets. Scale bars, 50 µm. Temporal kymographs in the bottom panels are generated along the white dashed line. Time scale bars, 20 s. (D) Representative image with eight color-coded neuronal spatial footprints. Scale bar, 50 µm. (E) Calcium activity traces extracted from the eight representative neurons shown in (D). NeuroPilot-recovered traces (blue) closely match the paired high-SNR reference traces (green) and recover calcium transients that are obscured in the raw low-SNR recordings (gray). Red boxes highlight representative calcium transients recovered by NeuroPilot. Scale bars, 2 ΔF/F and 20 s. (F) Pearson correlation coefficients between raw or NeuroPilot-recovered calcium traces and the paired high-SNR reference traces. (G) SNR of raw and NeuroPilot-recovered calcium traces. For (F) and (G), *P* < 0.0001, two-sided paired t-test; n = 115 neurons. Center lines indicate medians, box limits indicate the 25th and 75th percentiles, and whiskers indicate the data range. (H) Representative displacement vector fields estimated from the raw recording at two time points. Arrows indicate the direction of local tissue displacement, and colors indicate displacement magnitude. (I-K) NeuroPilot restores fine neuritic structures in a representative FOV. Representative images compare the raw low-SNR data (I), the NeuroPilot-restored result (J), and the paired high-SNR reference (K). Red boxes indicate zoomed regions shown in the adjacent insets, and temporal kymographs are generated along the white dashed line. Scale bars are as in (A–C). (L) Representative image with eight color-coded neuritic regions. Scale bar, 50 μm. (M) Calcium activity traces extracted from the eight representative neuritic regions shown in (L). Trace colors and red boxes are as in (E). Scale bars, 2 ΔF/F and 20 s. (N) Pearson correlation coefficients between raw or NeuroPilot-recovered traces and the paired high-SNR reference traces. (O) SNR of raw and NeuroPilot-recovered traces from the same neuritic regions. For (N) and (O), *P* < 0.0001, two-sided paired t-test; n = 143 neuritic regions. Box plot definitions are as in (F). (P) Representative displacement vector fields estimated from the neurite-enriched FOV at two time points. Arrows and color coding are as in (H). (Q-S) Time courses of x- and y-axis displacement in raw and NeuroPilot-corrected recordings. Data in (Q), (R), and (S) correspond to the fields of view shown in (A), (I), and **Figure S2**, respectively.

We next tested NeuroPilot in a neurite-enriched FOV, where photon noise and motion-induced deformation obscure thin neuronal processes. Compared with raw recordings, NeuroPilot-restored data showed more continuous neuritic morphology and more stable kymographs that better matched the paired high-SNR reference (Figures 2I-2K). Activity traces extracted from representative neuritic regions similarly showed closer agreement with reference traces after restoration (Figures 2L and 2M). Across 143 neuritic regions, NeuroPilot increased the Pearson correlation between extracted traces and the paired high-SNR reference from 0.280 ± 0.006 to 0.771 ± 0.007 (mean ± SEM; two-sided paired t-test, *P* < 0.0001; Figure 2N) and increased trace SNR from 1.58 ± 0.01 dB to 6.24 ± 0.08 dB (mean ± SEM; two-sided paired t-test, *P* < 0.0001; Figure 2O). Displacement vector fields and residual displacement traces showed non-uniform motion along both spatial axes and reduced frame-to-frame displacement after NeuroPilot correction (Figures 2P-2S).

We further evaluated NeuroPilot using an independent paired two-photon microscopy dataset, focusing on neuronal signal recovery across the imaged population (Figure S2). Suite2p^26,27^ detected only 35 neuronal footprints in the raw low-SNR recordings, whereas detection increased to 132 after NeuroPilot restoration, compared with 118 in the paired high-SNR reference recordings (Figure S2D; Methods). We assessed trace similarity and signal-to-background ratio relative to the paired reference. NeuroPilot also increased the Pearson correlation between extracted traces and the paired reference from raw correlation 0.2158 ± 0.0005 to NeuroPilot correlation 0.8656 ± 0.0007 and increased SBR from raw SBR 1.00640 ± 0.00004 dB to NeuroPilot SBR 1.50759 ± 0.00067 dB (two-sided paired t-test, *P* < 0.0001; Figure S2E). Representative traces showed reduced background fluctuations and clearer calcium transients after restoration, bringing extracted activity patterns closer to the paired reference recordings (Figure S2F).

### NeuroPilot links spinal neuronal activity across stimulus conditions and longitudinal sessions

We applied NeuroPilot to calcium imaging of the spinal cord in freely moving animals^33–35^, where limited photon collection, motion-induced global displacement, and muscle-driven local tissue deformation compromise image quality^7,36–38^. NeuroPilot expands the experimental capabilities of freely moving spinal cord imaging, from stable single-session readout to cross-condition and longitudinal analysis at single-neuron resolution. Using a custom grooved spinal window^39^, we applied this strategy to track spinal neuronal activity across different stimulus conditions and longitudinal imaging sessions (Figures 3A-3C; Methods).

**Figure 3.**
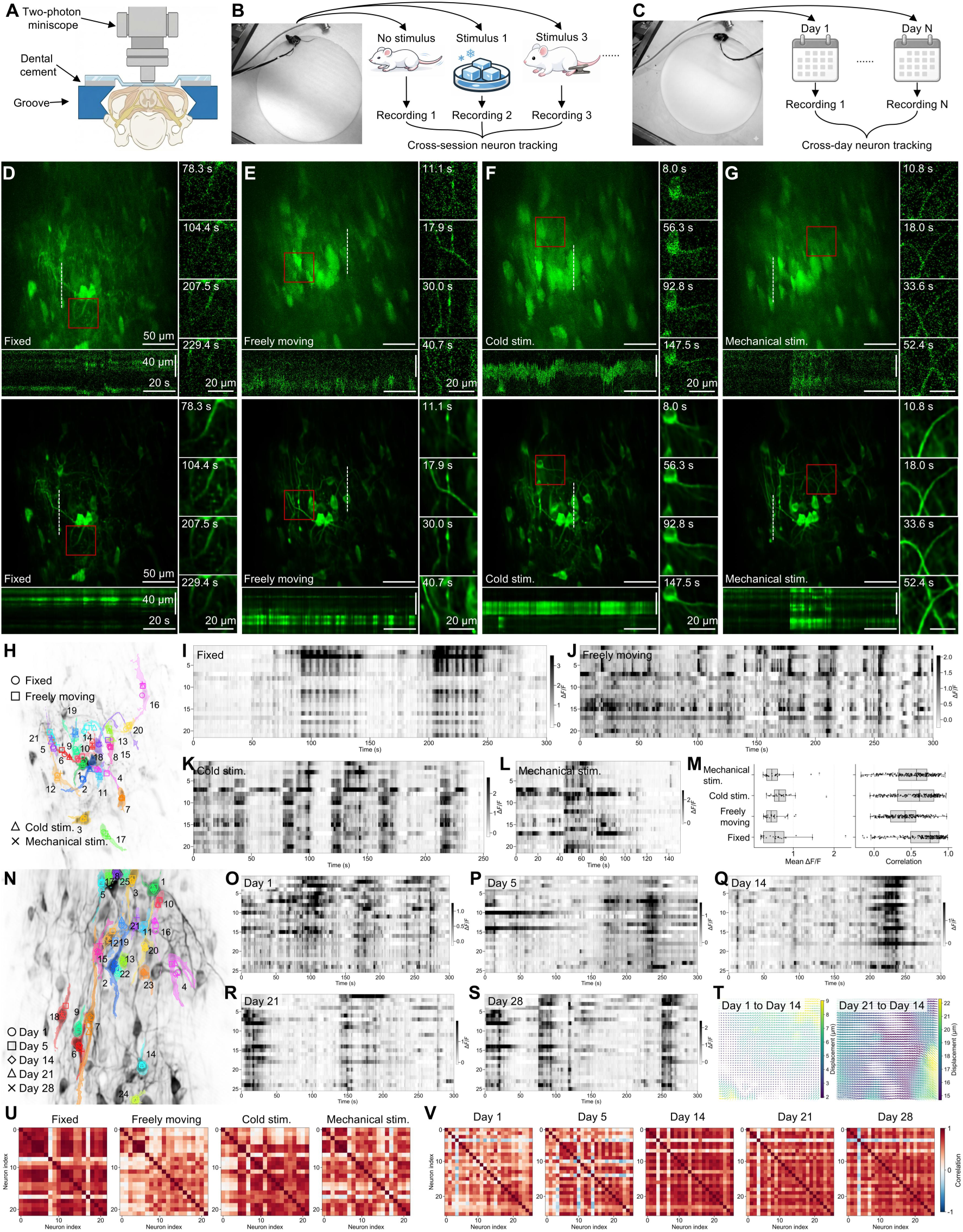
NeuroPilot enables cross-condition and longitudinal tracking of spinal cord neurons. (A) Schematic of the two-photon miniscope setup for spinal cord imaging in freely moving mice. The miniscope is positioned above a custom grooved spinal window. (B) Cross-session tracking of neuronal activity across different stimulus conditions within the same day. (C) Longitudinal tracking of matched neuronal populations across multiple days. (D-G) Example spinal cord imaging fields of view across four experimental conditions: spinal column fixation (D), freely moving behavior (E), cold stimulation (F), and hind-paw pinch (G). The upper and lower rows show MIPs from the raw and NeuroPilot-restored recordings, respectively. Red boxes indicate zoomed regions shown in the adjacent insets. Temporal kymographs were generated along the white dashed lines and are shown beneath each main image. Scale bars, 50 µm (main images), 20 µm (insets), and 40 µm by 20 s (kymographs). (H) Spatial map of neurons tracked across the four recording conditions. A total of 21 neurons were matched across all conditions. Marker shapes indicate recording condition, marker positions denote neuronal footprint centroids, and color-coded masks indicate the spatial footprints of individual neurons. (I-L) Calcium activity heatmaps of the 21 neurons tracked across all four recording conditions: spinal column-fixed preparation (I), freely moving imaging (J), cold stimulation (K), and nociceptive mechanical stimulation (L). Grayscale indicates ΔF/F amplitude. (M) Quantification of calcium activity across the four recording conditions. Left: distribution of mean ΔF/F values for the tracked neurons. Each dot represents one neuron (n = 21). Right: distribution of pairwise activity correlations within the tracked neuronal population. Each dot represents one neuron pair. Center lines indicate medians, box limits indicate the 25th and 75th percentiles, and whiskers indicate the data range. (N) Spatial map of longitudinally tracked neurons in mice after SNI. A total of 25 neurons were tracked across five imaging sessions on days 1, 5, 14, 21, and 28. Color-coded contours indicate individual neuronal footprints, and marker shapes indicate imaging day. The background image shows the standard-deviation projection from the Day 14 recording. (O-S) Calcium activity heatmaps of the 25 neurons tracked across all five longitudinal recording sessions: Day 1 (O), Day 5 (P), Day 14 (Q), Day 21 (R), and Day 28 (S). Grayscale indicates ΔF/F amplitude. (T) Representative cross-day displacement vector fields required to register the Day 1 and Day 21 recordings to the Day 14 reference. Arrows indicate local displacement direction, and colors indicate displacement magnitude in micrometers. (U-V) Pairwise activity correlation matrices for the cross-condition tracked neurons (U) and longitudinally tracked neurons (V). Colors indicate correlation coefficients.

We first examined recordings acquired from the same spinal FOV during spinal column fixation, freely moving behavior, cold stimulation, and nociceptive mechanical stimulation. Raw recordings showed condition-dependent motion and disrupted kymographs, whereas NeuroPilot processing improved spatial continuity and temporal stability across all four conditions (Figures 3D-3G and Video S2). This stabilization allowed 21 neurons to be matched across the four recording conditions in the same animal (Figure 3H; Methods). Because NeuroPilot linked the same 21 neurons across all four conditions, we compared activity changes within a shared spinal ensemble. The tracked-neuron heatmaps showed condition-associated differences in ΔF/F dynamics across the four experimental conditions (Figures 3I-3L)^40^. Quantification of mean activity, pairwise correlations, and correlation matrices further indicated that these conditions differed in both single-neuron activity levels and ensemble coordination (Figures 3M and 3U; Methods).

We extended NeuroPilot to longitudinal spinal cord imaging, enabling repeated recordings to be restored and integrated for cross-session neuronal identification and activity tracking. In a sciatic nerve injury (SNI) model, NeuroPilot linked 25 spinal cord neurons across five imaging sessions spanning day 1 before injury and days 5, 14, 21, and 28 after injury (Figures 3N-3S and Video S3; Methods). NeuroPilot corrected cross-day tissue deformation and aligned the longitudinal recordings into a shared reference frame (Figure 3T; Methods). Raw longitudinal recordings showed session-dependent structural instability and discontinuous kymographs, whereas NeuroPilot-restored recordings showed clearer cellular structure and more continuous spatiotemporal signals across the five SNI sessions (Figures S3A and S3B; Video S3). Frame-wise displacement analysis further showed that NeuroPilot reduced large motion excursions along the x and y axes, decreasing displacements of up to 30 μm in the raw recordings to near-zero residual values after correction (Figures S3C and S3D). Representative displacement fields captured the spatially heterogeneous motion present in individual sessions (Figure S3E). Pairwise activity correlation matrices from the longitudinally tracked neurons allowed activity patterns within the same spinal ensemble to be compared directly as they changed over days after injury (Figure 3V).

We further examined whether NeuroPilot could maintain neuronal identity across longitudinal spinal cord recordings acquired multiple days and sensory conditions. After registration, the same spinal FOV was aligned across five imaging days, yielding 96 tracked neuronal footprints (Figure S4A; Methods). This larger matched population allowed activity dynamics to be followed across spinal fixation, freely moving behavior, cold stimulation, hind-paw pinch, and air puff stimulation over time (Figure S4B and Video S2). Pairwise correlation matrices showed that ensemble organization varied across day-condition combinations, with sensory stimulation producing more structured population-wide correlations than the fixed baseline in several sessions (Figure S4C). Quantification of mean ΔF/F and pairwise activity correlation further showed that spinal activity was not uniform across the longitudinal series: activity levels varied across days and conditions, whereas population coordination was most strongly organized by stimulus context (Figures S4D and S4E). Grouping by day and stimulus condition clarified these patterns: mean activity varied modestly across days, whereas stimulus-dependent differences were more pronounced in both activity level and pairwise correlation (Figures S4F-S4I).

### NeuroPilot supports neural activity readout across imaging modalities and spatial scales

Different imaging modalities offer complementary strengths for neural activity readout, with three-photon microscopy enabling deep-tissue imaging^41^ and one-photon imaging supporting large-field-of-view recordings^8,19,20,22^. We applied NeuroPilot to head-mounted three-photon calcium imaging extending through the cortex and into hippocampal CA1 (Figure 4A and Video S4)^11,42^. In volumetric recordings acquired to approximately 1,000 μm, signal quality deteriorated progressively with depth, whereas NeuroPilot improved cellular separation across cortical layers, the corpus callosum, and CA1 (Figure 4B). Representative planes at 495 μm, 670 μm, and 908 μm showed improved neuronal visibility from L4/5 through L6 to hippocampal CA1 (Figure 4C). Restored recordings yielded cleaner calcium traces with more clearly resolved transients and supported segmentation of neuronal footprints at each depth (Figures 4D and 4E). This improvement was maintained during a 100-min three-photon CA1 recording. NeuroPilot increased cellular contrast and restored continuous spatiotemporal structure throughout the session (Figure 4F), while correcting motion excursions of up to 25 μm along both spatial axes (Figure 4G). Population heatmaps and representative traces showed reduced noise while preserving calcium transients throughout the recording, extending deep-tissue restoration to sustained population imaging (Figure 4H). We next tested NeuroPilot on head-mounted one-photon miniscope recordings of the prefrontal cortex (PFC)^6,31,43^. Despite broad background fluorescence and fixed-pattern noise, NeuroPilot improved cellular separability across the FOV and supported large-scale neuronal segmentation (Figures 4I and 4J). Extracted traces showed reduced motion-associated fluctuations, while kymographs showed improved spatial continuity and reduced motion and fixed-pattern artifacts (Figures 4K and 4L).

**Figure 4.**
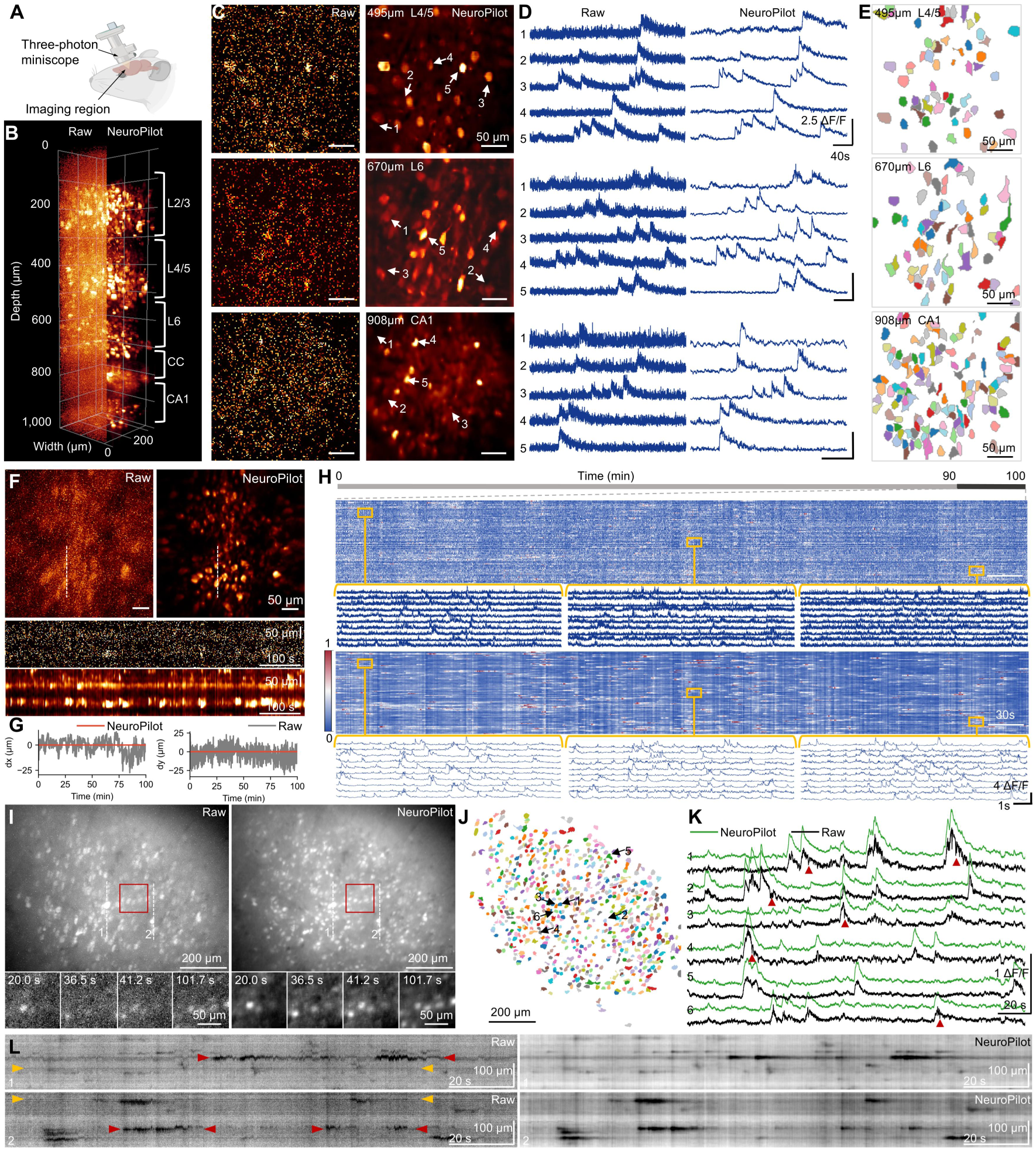
NeuroPilot generalizes across three-photon and one-photon calcium imaging. (A) Schematic of the three-photon miniscope configuration for in vivo deep-brain imaging. (B) Volumetric rendering of raw (left) and NeuroPilot-restored (right) three-photon imaging data acquired to a depth of 1,000 µm. Brackets indicate cortical layers and anatomical landmarks, including L2/3, L4/5, L6, corpus callosum (CC), and hippocampal CA1. (C) Representative imaging planes comparing raw (left) and NeuroPilot-restored (right) three-photon recordings at 495 µm in L4/5, 670 µm in L6, and 908 µm in CA1. Scale bars, 50 µm. (D) Calcium activity traces extracted from the numbered neurons shown in (C). Left: traces extracted from raw recordings; right: corresponding traces from NeuroPilot-restored data. Scale bars, 2.5 ΔF/F and 40 s. (E) Segmentation maps of neuronal spatial footprints extracted from NeuroPilot-restored recordings. Color-coded regions indicate individual neuronal footprints. Scale bars, 50 µm. (F) NeuroPilot restoration of a 100-min three-photon CA1 recording. MIPs of raw and NeuroPilot-restored data are shown at the top, with temporal kymographs generated along the dashed lines shown below. Scale bars, 50 μm (images) and 50 μm by 100 s (kymographs). (G) Time courses of x- and y-axis displacement over 100 min before and after NeuroPilot motion correction. Gray traces indicate raw recordings, and red traces indicate NeuroPilot-corrected recordings. (H) Population activity heatmaps and representative calcium traces from the same 100-min session. Raw data are shown at the top, and NeuroPilot-restored data are shown at the bottom. Enlarged trace windows indicated by yellow boxes show reduced noise while preserving calcium transients across the recording. Scale bars, 30 s (heatmaps) and 1 s by 4 ΔF/F (traces). (I) Comparison of raw and NeuroPilot-restored one-photon miniscope imaging. Top panels show full-field MIPs, and bottom panels show magnified snapshots of the red-boxed region. Scale bars, 200 μm (top) and 50 μm (bottom insets). (J) Segmentation map of neuronal spatial footprints extracted from NeuroPilot-restored data. Color-coded masks indicate individual ROIs, and numbered arrows indicate representative neurons. Scale bar, 200 μm. (K) Calcium activity traces from the six representative neurons shown in (J). Raw traces (black) are plotted with corresponding NeuroPilot-restored traces (green). Red arrowheads indicate motion-induced artifacts in the raw traces that are reduced after NeuroPilot restoration. Scale bars, 1 ΔF/F and 20 s. (L) Kymographs comparing raw (left) and NeuroPilot-restored (right) signals along the dashed lines shown in (I). Colored arrowheads indicate examples of reduced motion artifacts (red) and fixed-pattern noise (yellow). Scale bars, 100 µm and 20 s.

NeuroPilot further extended across fluorescence channels and spatial scales. In dual-color two-photon imaging of vS1^44^, processing improved the spatial and temporal definition of both GCaMP6s and mCherry signals, supporting neuronal segmentation and trace extraction from GCaMP6s channel (Figures S5A-S5C and Video S5). Channel-wise correlation analysis further assessed signal consistency in the two fluorescence channels (Figure S5D). NeuroPilot also recovered local structures and spatiotemporal continuity across a millimeter-scale cortical FOV spanning 6 mm^45^, extending the workflow from cellular fields of view to large-area imaging (Figure S5E and Video S4).

### NeuroPilot enables robust signal recovery across deep-brain and subcellular imaging targets

We examined whether NeuroPilot could recover neuronal activity from miniature two-photon recordings acquired in deep brain regions. In the lateral entorhinal cortex (LEC, DV = 4.30 mm), miniature two-photon recordings acquired through a GRIN lens showed that NeuroPilot improved visualization of neuronal morphology and supported segmentation of neuronal footprints that were poorly delineated in the raw recording (Figures 5A-5C and Video S4). Extracted traces showed reduced noise and more clearly resolved calcium transients (Figure 5D), and motion trajectories, displacement fields, and kymographs confirmed improved spatial stability after correction (Figures 5E-5H). FRC analysis^46^ showed that the estimated spatial resolution improved from 5.784 μm in the raw recording to 2.008 μm after restoration, while SBR increased from 2.4266 dB to 13.5691 dB (two-sided paired t-test, *P* < 0.0001; Figures 5I and 5J; Methods). The resulting neuronal traces supported temporal similarity analysis and low-dimensional visualization of population activity states (Figures 5K and 5L). We also evaluated NeuroPilot in miniature two-photon recordings acquired through a GRIN lens in hippocampal CA1 (DV = 1.20 mm; Figure 5M and Video S4). NeuroPilot improved neuronal visibility and increased the number of segmented spatial footprints by Suite2p from 52 in the raw recording to 414 after restoration (Figures 5N and 5O). Representative traces showed reduced noise and clearer calcium events (Figure 5P). FRC analysis further showed that the estimated spatial resolution improved from 16.395 μm in the raw recording to 2.239 μm after restoration, while SBR increased from 0.716 dB to 12.360 dB (two-sided paired t-test, *P* < 0.0001; Figures 5Q and 5R). The restored population signals further supported similarity analysis and visualization of activity trajectories over the recording session (Figures 5S and 5T).

**Figure 5.**
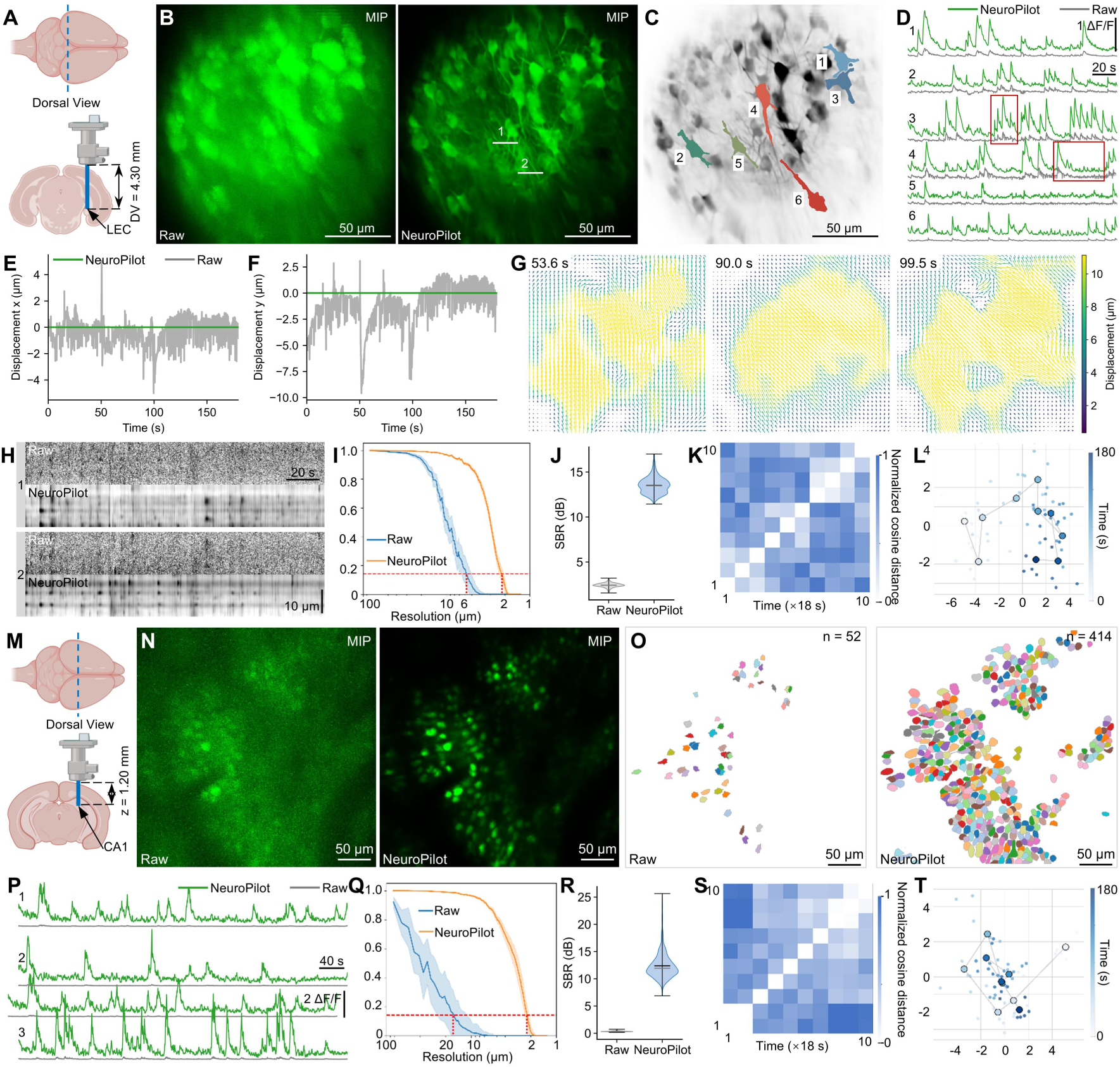
NeuroPilot enables high-fidelity calcium imaging in deep brain regions. (A) Deep-brain two-photon miniscope imaging of the LEC in freely moving mice at DV = 4.30 mm. The coronal section shows the lens implantation trajectory corresponding to the dashed line in the dorsal view. (B) Raw and NeuroPilot-restored MIPs from the LEC. Numbered white lines indicate sampling lines for kymograph analysis. Scale bars, 50 µm. (C) Segmentation map of neuronal spatial footprints extracted from NeuroPilot-restored LEC data. Color-coded masks indicate individual ROIs. Scale bars, 50 µm. (D) Calcium activity traces from neurons shown in (C). Raw traces are shown in gray and NeuroPilot-restored traces in green. Red boxes highlight representative recovered calcium transients. Scale bars, 1 ΔF/F and 20 s. (E-F) Time courses of x-and y-axis displacement before and after NeuroPilot motion correction. (G) Representative displacement vector fields at selected time points. Arrows indicate local displacement direction, and colors indicate displacement magnitude in micrometers. (H) Spatiotemporal kymographs along lines 1 and 2 shown in (B). Scale bars, 20 s and 10 µm. (I) Spatial resolution estimated by FRC. Solid lines and shaded bands indicate mean ± s.d. for raw and NeuroPilot-restored recordings. Red dotted lines indicate resolution estimates at the 1/7 criterion. (J) SBR of calcium traces extracted from the same neuronal regions in raw and NeuroPilot-restored recordings. (K) Temporal similarity matrix of population activity. Each entry represents the normalized cosine similarity between activity vectors across 18-s time bins; the 10 × 10 matrix spans the 180-s recording. (L) Two-dimensional visualization of neural population activity trajectories. Points are color-coded by time to show the temporal progression of population activity states. (M) Schematic of miniscope imaging targeting CA1 at DV = 1.20 mm. (N) Raw and NeuroPilot-restored MIPs from hippocampal CA1. Scale bars, 50 µm. (O) Segmentation maps of neuronal spatial footprints extracted from raw (left, n = 52) and NeuroPilot-restored (right, n = 414) CA1 data. Scale bar, 50 µm. (P) Example calcium activity traces from hippocampal CA1 neurons. Raw traces are shown in gray and NeuroPilot-restored traces in green. Scale bars, 2 ΔF/F and 40 s. (Q) Spatial resolution estimated by FRC for raw and NeuroPilot-restored CA1 images. (R) SBR of neuronal signals extracted from raw and NeuroPilot-restored CA1 data. (S) Population activity similarity matrix computed from normalized cosine similarity across 18-s time bins. (T) Low-dimensional projection of neural population activity trajectories, color-coded by session time.

To assess its applicability beyond neuronal imaging, we applied NeuroPilot to vascular structures and mitochondrial calcium imaging. In spinal vascular imaging of freely moving animals, NeuroPilot improved the continuity of tubular structures and stabilized their spatiotemporal profiles (Figure S5F and Video S5). In mitochondrial calcium imaging, processing recovered more continuous mitochondrial structures and local intensity profiles (Figure S5G), increased local spatial correlation (Figure S5H), and supported mitochondrial segmentation and skeletonization using the MINA pipeline (mitochondrial network analysis, Figure S5I)^47^. Spatiotemporal profiles showed reduced motion-associated discontinuities (Figure S5J), enabling quantitative analysis of mitochondrial footprint, network branching, and branch length (Figure S5K).

### NeuroPilot reveals behavioral-state-dependent mPFC activity across 24-hour recordings

We evaluated NeuroPilot for continuous population-level analysis during spontaneous behavior across a full circadian cycle. Continuous 24-hour neural imaging provides a unique opportunity to examine how population activity evolves across spontaneous behavioral transitions, sleep-wake cycles, and circadian time^48–51^. We applied NeuroPilot to head-mounted two-photon calcium recordings from the medial prefrontal cortex (mPFC) of freely behaving mice, aligned with multimodal behavioral and physiological annotations spanning 12 states over 24 hours (Figure S6 and Video S6)^52^. Across three animals, the recordings captured diverse sleep and behavioral repertoires whose prevalence varied across Zeitgeber time (Figure S7A). A representative 24-hour recording from Mouse 2 captured repeated transitions among REM sleep, non-REM sleep, microarousals, locomotion, grooming, nesting, and other spontaneous behaviors (Figures 6A and 6B; Video S6). NeuroPilot enhanced the clarity of neuronal structures across the FOV, while kymographs showed more continuous spatiotemporal structure throughout the recording (Figures 6C and 6D). Behavior-aligned traces further showed reduced noise and more clearly resolved calcium transients across active and inactive states (Figure 6E).

**Figure 6.**
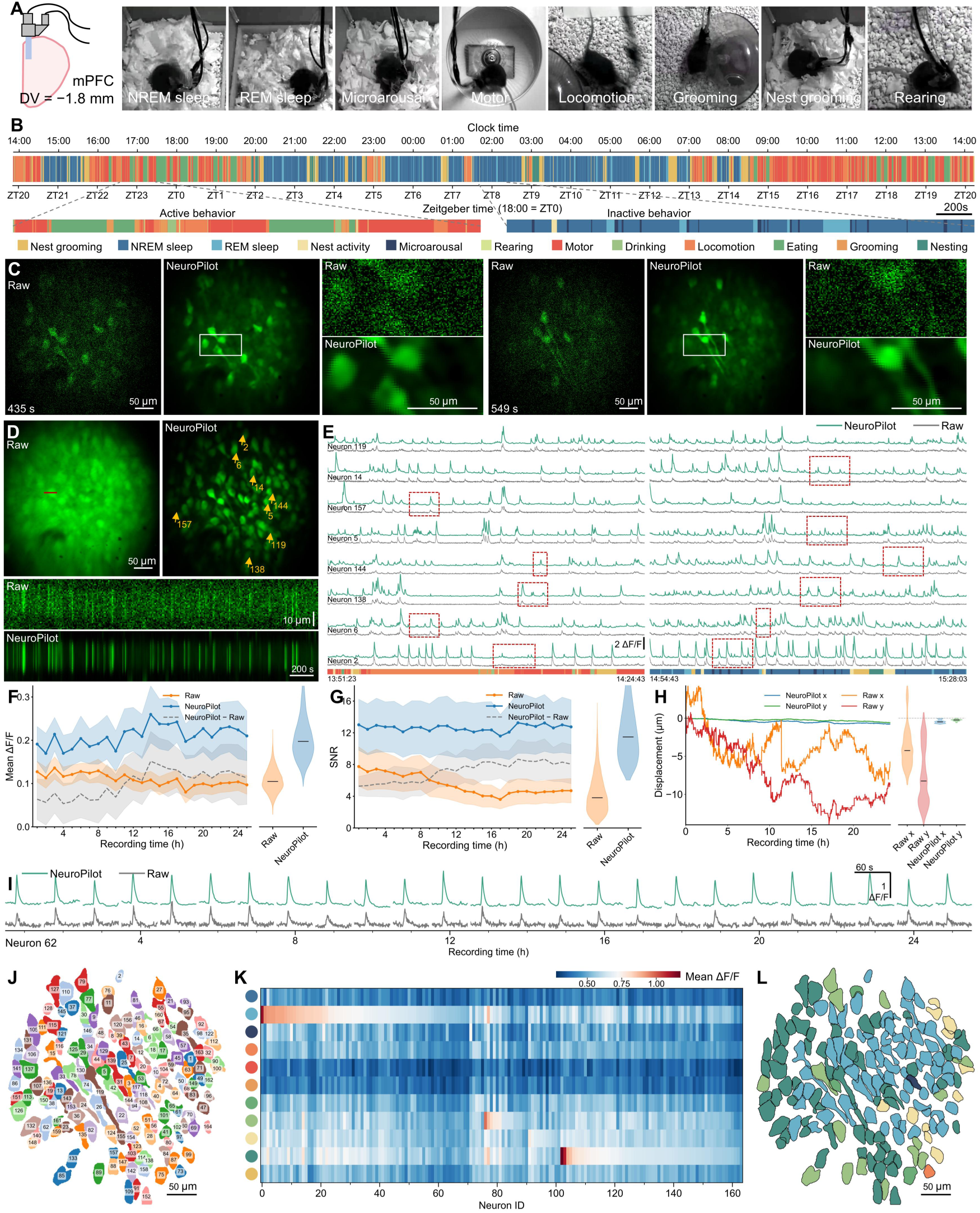
NeuroPilot sustains stable mPFC neuronal imaging across a 24-hour sleep–wake cycle. (A) Schematic of the mPFC imaging site at DV = −1.8 mm and example video frames showing representative behavioral states observed during the 24-hour recording. (B) Behavioral ethogram from the continuous 24-hour recording. Colors indicate 12 annotated behavioral states plotted against clock time and Zeitgeber time (18:00 = ZT0). Lower panels show magnified examples of active and inactive periods. Scale bar, 200 s. (C) Raw and NeuroPilot-restored calcium imaging data at representative time points. White boxes indicate regions shown at higher magnification in the right panels. Scale bars, 50 μm. (D) STD projections (top) and kymographs (bottom) comparing raw and NeuroPilot-restored data. Kymographs are generated along the red sampling line. Scale bars, 50 μm (top) and 10 μm by 200 s (bottom). (E) Calcium activity traces from neurons shown in (D). Raw traces are shown in gray and NeuroPilot-restored traces in green. Red dashed boxes highlight several representative calcium transients. Color-coded bars indicate behavioral states during the corresponding recording periods. Scale bar, 2 ΔF/F. (F-G) Mean ΔF/F (F) and SNR (G) of mPFC population activity over the 24-hour recording. Raw data are shown in orange and NeuroPilot-restored data in blue; dashed gray traces indicate NeuroPilot minus raw values. Shaded areas indicate ± s.e.m. across neurons. (H) Cumulative x- and y-axis displacement over the 24-hour recording. (I) Example long-duration calcium trace from neuron 62 showing raw and NeuroPilot-restored activity over the recording. Scale bars, 60 s and 1 ΔF/F. (J) Individual neurons (n = 164) are shown in distinct colors, and numbers indicate neuron IDs. Scale bar, 50 μm. (K) Heatmap of mean ΔF/F across behavioral states for all segmented neurons. Behavioral states are color-coded as in (B), and neurons are ordered by similarity of behavioral activity profiles. (L) Anatomical projection of the behavioral activity groups shown in (K). Neuronal footprints are color-coded by the behavioral state with the highest mean activity. Scale bar, 50 μm.

Across the 24-hour recording, NeuroPilot increased the mean population ΔF/F from 0.110 ± 0.002 to 0.210 ± 0.003 and increased SNR from 5.45 ± 0.13 dB to 12.61 ± 0.25 dB (mean ± SEM; two-sided paired t-test, *P* < 0.0001; n = 164 neurons; Figures 6F and 6G; Methods). In addition, the raw recording showed a progressive photobleaching-associated decline in SNR, which was corrected after NeuroPilot processing. Raw recordings exhibited cumulative displacements of up to 15 μm, while NeuroPilot reduced residual displacement to 0.5 μm along the x axis and 0.5 μm along the y axis (Figure 6H). This combined signal and spatial stabilization supported continuous extraction of 164 neuronal footprints and preserved single-neuron traces across the full recording (Figures 6I and 6J; Methods). Representative neurons remained detectable across temporally separated behavioral epochs, enabling comprehensive characterization of their activity patterns across behavioral states over 24 hours (Figure S6).

With a stable 164-neuron population established across the full recording, we characterized how mPFC activity was organized across spontaneous behavioral states^53,54^. State-averaged activity profiles identified distinct neuronal ensembles associated with sleep, wakefulness, and active behaviors, including a prominent group with elevated activity during REM sleep (Figure 6K)^55^. When mapped back onto the imaging field, state-associated neurons were spatially intermingled rather than confined to discrete anatomical domains, suggesting that behavioral-state-associated activity was broadly distributed across the recorded mPFC population (Figure 6L).

This state dependence also emerged at the level of population coordination. Across three mice, pairwise correlations varied systematically among behavioral states, with NREM sleep exhibiting stronger ensemble synchrony than REM sleep and several active states (Figure S7B). Continuous correlation trajectories changed alongside spontaneous behavioral transitions, indicating that mPFC population coordination varied in concert with transitions among sleep, quiet wakefulness, and active behavior (Figure S7C). Two additional animals, containing 171 and 161 neurons, showed similarly heterogeneous state-associated activity profiles and spatially distributed functional ensembles (Figures S7D-S7I). These analyses show that NeuroPilot enables day-long tracking of mPFC activity and ensemble coordination across natural sleep-wake and behavioral states.

### mPFC population activity distinguishes behavioral states and reorganizes across state transitions

Having established stable population readout across 24 hours, we assessed how strongly mPFC activity reflected ongoing behavioral states. Population-based classifiers distinguished sleep from wakefulness and separated four major behavioral categories, while decoding of the full 12-state ethogram revealed a structured pattern of errors concentrated among behaviorally related states (Figures 7A-7D). Consistent with this organization, hierarchical clustering grouped states with similar population activity patterns (Figure 7E).

**Figure 7.**
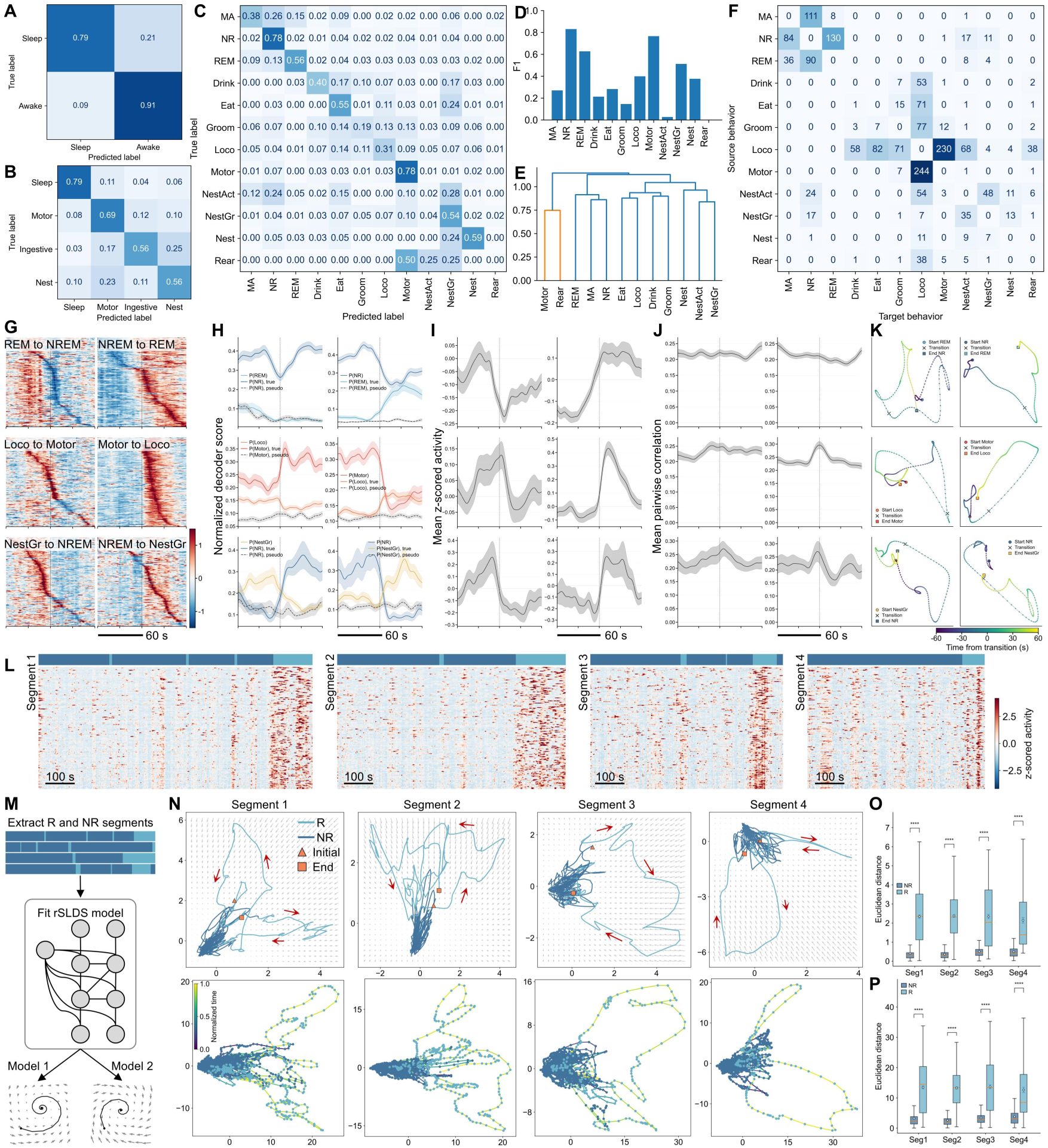
24-hour mPFC population activity encodes spontaneous behavioral states and transitions. (A-C) Confusion matrices for decoding behavioral labels from mPFC population activity. Matrices show binary sleep-wake decoding (A), decoding of four major behavioral categories (B), and decoding of 12 fine-grained behavioral labels (C). Values indicate the fraction of time points assigned to each predicted label for each true label. (D) Per-behavior F1 scores from the 12-class behavioral decoder. (E) Dendrogram of the 12 behavioral states based on pairwise similarity of mPFC population activity patterns. (F) Source-to-target behavioral transition matrix. Rows indicate source behaviors, columns indicate target behaviors, and values indicate transition frequency. (G) Transition-aligned heatmaps of single-neuron activity around representative behavioral transitions: REM-to-NREM (n = 83), NREM-to-REM (n = 123), locomotion-to-motor (n = 89), motor-to-locomotion (n = 100), nest grooming-to-NREM (n = 17), and NREM-to-nest grooming (n = 11). Neurons are ordered by their transition-associated response profiles. Scale bar, 60 s. (H) Transition-aligned decoder-score trajectories for selected state pairs. Solid traces indicate true transition-aligned responses, and dashed traces indicate pseudo-transition controls. Shading indicates ± s.e.m. across transition events. Scale bar, 60 s. (I) Mean z-scored population activity aligned to the same behavioral transitions as in (G). Scale bar, 60 s. (J) Mean pairwise neuronal correlation aligned to behavioral transitions. Scale bar, 60 s. (K) Representative low-dimensional neural trajectories around behavioral transitions. Markers indicate transition time. (L) Z-scored mPFC population activity across continuous NREM-REM transition segments. Aligned sleep-state annotations are shown above. Scale bars, 100 s. (M) rSLDS analysis workflow. REM and NREM segments are extracted to fit an rSLDS model and infer latent dynamics. (N) Representative rSLDS latent trajectories and PCA trajectories. Top, rSLDS trajectories overlaid on inferred local vector fields; arrows indicate local flow direction. Bottom, PCA trajectories computed from neural activity. Colors indicate sleep state or normalized time as shown. (O-P) Distances to the NREM centroid in rSLDS latent space (O) and PCA space (P) across the same four representative segments as in (N). Distances are shown separately for NREM and REM epochs. Center lines indicate medians, boxes indicate the 25th and 75th percentiles, whiskers indicate the data range, and white circles indicate means. *P* < 0.0001, two-sided paired t-test.

We next examined how this representation evolved as animals moved between states. The 24-hour recordings contained repeated transitions among sleep, locomotor, motor, and nesting-related behaviors (Figure 7F)^56^. Aligning neuronal activity to six frequently observed transitions revealed transition-specific ensembles whose activity changed before and after state boundaries (Figure 7G). Behavioral decoder scores shifted around the true transitions but not around matched pseudo-transition events, indicating that these changes were temporally associated with behavioral switching rather than nonspecific fluctuations (Figure 7H). Transitions were accompanied by transition-specific changes in population-averaged activity and pairwise neuronal correlation, with the direction, magnitude, and temporal profile of these changes varying across sleep, motor, and nesting-related transitions (Figures 7I and 7J). Low-dimensional trajectories followed transition-specific paths, suggesting history-dependent population dynamics (Figure 7K)^57^.

We then focused on the recurrent transition between NREM and REM sleep^58^. Continuous segments showed coordinated evolution of neuronal activity as the population moved between the two sleep states (Figure 7L and Video S7). We fitted a recurrent switching linear dynamical system to these segments to infer state-dependent latent trajectories and local vector fields (Figure 7M and Video S7; Methods)^59–63^. In both the rSLDS representation and a model-independent PCA projection, NREM activity occupied a relatively confined region, whereas REM trajectories extended farther from the NREM population centroid (Figure 7N; Methods). Across four representative segments, distances from the NREM centroid were greater during REM than during NREM in both latent spaces (two-sided paired t-test, *P* < 0.0001; Figures 7O and 7P; Methods). This analysis suggested an attractor-like organization in latent space, with NREM trajectories clustered around a common region and REM states located farther away.

The same analyses in two additional mice reproduced the principal findings. Twelve-state decoders recovered behavioral information with state-dependent F1 scores (Figures S8A, S8B, S8J, and S8K). Transition-aligned activity, decoder evidence, population correlation, and low-dimensional trajectories also varied around spontaneous behavioral transitions (Figures S8E-S8I and S8N-S8R). REM activity also remained farther from the NREM centroid, with corresponding differences in model-inferred latent trajectories across representative sleep segments (Figures S8C, S8D, S8L, and S8M).

## Discussion

Advances in neural imaging increasingly support large-scale and long-duration recordings across anatomical depths, behavioral contexts, and imaging modalities. Realizing the scientific value of these advances requires converting heterogeneous recordings into stable and interpretable representations of neural activity. NeuroPilot addresses this challenge through an agentic framework that integrates a multimodal large language model with modular computational imaging tools to adaptively organize analysis from raw recordings to population-level readouts. Its central contribution lies in orchestrating restoration, non-rigid registration, segmentation, tracking, quality assessment, and downstream analysis within an iterative, context-aware, and traceable workflow. In this formulation, intermediate outputs, including restored stacks, deformation fields and quality-control metrics, are recorded as quality-control evidence and used to guide subsequent processing, review, and parameter refinement.

In paired low- and high-SNR two-photon recordings, NeuroPilot-restored calcium traces more closely matched simultaneously acquired reference signals, showing that improved image quality translated into more faithful functional readout. Across one-, two-, and three-photon calcium imaging, the framework supported stable signal extraction from deep-brain, large-field, volumetric, multiplexed, and freely-moving miniature-microscope recordings^64–66^. Its application to neurites, vascular structures, and mitochondrial calcium signals further indicates that a shared agentic workflow can accommodate biological targets with distinct spatial organization and degradation profiles without requiring a separately constructed end-to-end pipeline for each dataset. In freely moving spinal cord imaging, NeuroPilot stabilized recordings across motion regimes, sensory conditions, and longitudinal sessions, enabling the same neurons to be tracked within shared reference frames. This capability revealed stimulus- and injury-associated changes in neuronal activity and ensemble coordination that would otherwise be obscured by tissue deformation and cross-session instability. In day-long mPFC imaging, NeuroPilot preserved stable population readout across natural sleep-wake and behavioral cycles, supporting analysis of distributed state-associated ensembles, state-dependent coordination, and transition-specific population trajectories. These analyses further distinguished NREM and REM dynamics, with NREM occupying a more confined latent region and REM extending into more dispersed population states.

The broader implication is that the future of neural imaging will depend not only on acquiring more data, but on building computational systems capable of turning those data into biological understanding. As experiments expand across scales, modalities, brain regions, and timescales, neuroscience is producing datasets whose volume and heterogeneity exceed what can be reliably interpreted through manually assembled workflows. Agentic infrastructure offers a path beyond this bottleneck by continuously adapting analytical strategies, documenting computational decisions, and integrating intermediate outputs into coherent population-level representations^67–69^. NeuroPilot illustrates how such a system can function as a computational bridge from measurement to meaning, transforming heterogeneous recordings into reproducible descriptions of neural activity, state, and dynamics. As neural datasets continue to grow in scale and complexity, this type of agentic infrastructure may become a necessary foundation for extracting biological insight from modern experiments.

A key next step is to develop NeuroPilot into an uncertainty-aware infrastructure for longitudinal neuronal surveillance. Future systems should not only recover activity from individual recordings, but maintain persistent, confidence-weighted representations of the same neurons across time, modalities, and experimental conditions. This requires explicit uncertainty estimates for deformation fields, neuronal footprints, calcium traces, and cross-session identities, enabling adaptive tool selection, parameter refinement, and human review before local errors propagate into downstream biological claims. At a broader level, these representations could populate a neuroscience data matrix in which each neuron is indexed simultaneously by functional dynamics, behavioral context, transcriptomic identity, morphology, projection pattern, anatomical position, electrophysiology, hemodynamics, and perturbation response. Such a matrix would convert fragmented measurements into an integrated description of neuronal identity and circuit function, allowing agentic systems to reason across modalities rather than process them in isolation. Benchmark-driven validation and cross-laboratory testing will be essential for establishing this framework as a scalable foundation for circuit-level inference and longitudinal neuroscience^70–73^.

A complementary direction is to move beyond in silico agents toward wet-lab experiments with microscopy in the loop. In this setting, the agent would not simply analyze completed recordings, but operate as an autonomous experimental partner that observes, reasons, and acts. Real-time estimates of motion, photobleaching, signal quality, tracking confidence, and neural state could guide focus, illumination, sampling, field-of-view selection, and perturbation timing as the experiment unfolds. NeuroPilot therefore points toward a robotic measurement architecture in which the microscope, analytical tools, and experimental interventions are coordinated through a continuous perception-decision-action loop. Such systems could make neural imaging adaptive rather than predetermined, maintaining the quality and interpretability of measurements while responding to biological dynamics in real time^74–76^.

## Limitations of the study

Several limitations define the current scope of NeuroPilot. First, although we developed dedicated modules for neural image restoration, non-rigid registration, and longitudinal tracking, the agentic capabilities of NeuroPilot remain bounded by the computational tools available in its library. The system can select, configure, and iteratively coordinate these modules on the basis of intermediate results, but it cannot yet create a new computational operator when existing tools are inadequate for an unfamiliar failure mode or imaging condition. A major next step will therefore be to extend NeuroPilot from tool orchestration to tool generation, enabling the language model to write, test, validate, and integrate new analysis modules in response to previously unseen data. Second, the current implementation operates primarily as an offline analysis system and does not yet support real-time processing during image acquisition. The computational demands of restoration, dense motion estimation, quality assessment, and iterative workflow refinement currently limit their use for low-latency experimental control. Developing a real-time implementation will require accelerated models, streaming data processing, hardware-aware scheduling, and rapid uncertainty estimation. Such advances could allow NeuroPilot to use incoming data to adjust focus, illumination, sampling rate, field-of-view selection, or perturbation timing, thereby linking analysis directly to microscopy acquisition and experimental intervention.

## Methods

### Image restoration and denoising module

To restore neural imaging data degraded by noise and low signal-to-noise ratio, we developed a self-supervised image restoration module based on a three-dimensional U-Net architecture, trained directly on raw recordings without the need for clean ground-truth data.

**Network architecture** The denoising model was implemented using a 3D U-Net architecture^77^ operating on spatiotemporal image volumes. The network consists of a multi-level encoder–decoder structure with skip connections that preserve high-resolution spatial information while capturing hierarchical features. The encoder path is composed of sequential convolutional blocks followed by downsampling operations, progressively increasing the number of feature channels. Each block consists of two consecutive 3D convolution layers with non-linear activation (ReLU). The decoder path performs symmetric upsampling and feature fusion by concatenating encoder features with upsampled representations, followed by convolutional refinement. A final 1×1×1 convolution maps the features to the restored output volume. All convolutions operate in three dimensions (time × height × width), enabling the model to jointly exploit spatial and temporal redundancy in calcium imaging data.

**Self-supervised training strategy** The model was trained in a self-supervised manner using only noisy recordings, without requiring paired clean targets. To construct training pairs, input patches were generated from raw data using structured masking strategies that exploit redundancy in space and time. Specifically, multiple sampling modes were implemented:

Spatial splitting (XY mode): input and target patches were constructed by partitioning pixels along spatial dimensions (e.g., interleaved rows or columns).

Temporal splitting (T mode): input and target patches were constructed from alternating frames along the temporal dimension.

Pixel dropout (Noise2Void-style^78^): a subset of pixels was randomly masked in the input, while the original signal served as the training target.

These sampling strategies enable self-supervised denoising by requiring the network to reconstruct withheld information from neighboring spatial or temporal observations. The sampling scheme was selected according to the form of redundancy exploited at each processing stage: XY interleaving was used to recover spatial cellular structure, temporal interleaving supported refinement from adjacent frames, and Noise2Void-style random pixel masking provided an alternative when interleaving along either dimension was unsuitable. The model was optimized with Adam using a combined L1 and L2 reconstruction objective.

**Inference and volumetric reconstruction** During inference, full image stacks were processed using a patch-based strategy to accommodate large volumes. Input volumes were divided into overlapping 3D patches, and each patch was independently processed by the trained network. The denoised patches were then merged to reconstruct the full volume, with overlap regions ensuring smooth transitions and reducing boundary artifacts. To ensure compatibility with the network architecture, input patches were padded to match required dimensions and cropped back to the original size after processing.

### Multi-scale non-rigid motion correction with PyLoReg-Net

To correct rigid and non-rigid motion in neural imaging recordings, we developed PyLoReg-Net, a coarse-to-fine optical-flow framework combining robust template construction, translation correction, hierarchical image encoding, local correlation matching, convolutional flow refinement, and iterative flow composition. PyLoReg-Net operates on single-channel image sequences and estimates a dense two-dimensional displacement field for every frame. The registration workflow consisted of an initial translation-only correction followed by two rounds of neural-network-based dense registration. During iterative refinement, the processed image stack was updated after each round to improve template estimation, whereas the original raw stack was retained without interpolation and warped only once using the final composed deformation field.

**Input normalization and spatial padding** The processed image stack and the corresponding raw stack were independently converted to 32-bit floating-point arrays and min-max normalized to the range [0,1]. Before neural-network inference, images were padded along the bottom and right boundaries using reflection padding so that both spatial dimensions were divisible by 16. Predicted flow fields were cropped back to the original image dimensions before application.

**Robust template construction** A robust global template was regenerated at every registration round from the current version of the processed image stack. Up to 2,000 frames were sampled uniformly across the recording. When the recording contained fewer than 2,000 frames, all available frames were included in the candidate pool. For computationally efficient template scoring, a centered 256×256 pixel region was extracted from each candidate frame and spatially sampled with a stride of four. Each resulting frame vector was mean-centered, standardized by its standard deviation, and normalized to unit _2_ norm. A consensus representation was calculated as the mean of the normalized feature vectors and was itself normalized to unit length. The similarity score for each candidate frame was then calculated as its dot product with the consensus vector, equivalent to cosine similarity after normalization. Frames were ranked by their consensus-similarity scores, and the highest-scoring 20% were selected, with a minimum of 50 frames retained whenever available. The full-resolution registration template was calculated as the pixel-wise mean of these selected frames. Template scoring was performed in batches of 256 frames on the CPU, with a numerical stabilization constant of 10^−6^. The selected frame indices, similarity scores, and template images were saved for quality control.

**Initial translation correction** The first registration round corrected global translational motion using phase correlation. Frames were processed sequentially, with each frame initially aligned to the preceding corrected frame. To limit cumulative drift, registration was re-anchored to the robust global template every 50 frames. For each image pair, the moving and reference images were mean-centered and transformed into the Fourier domain. Their normalized cross-power spectrum was calculated, and its inverse Fourier transform produced a phase-correlation map. The location of the maximum correlation response was used to estimate the horizontal and vertical translation. A numerical stabilization constant of 10^−9^ was included during cross-power normalization. The maximum accepted translation magnitude was set to 20% of the shorter image dimension. Estimates exceeding this threshold were rejected and replaced with zero displacement. Following alignment to the preceding corrected frame, normalized cross-correlation was calculated between the registered frame and its reference. If the correlation was below 0.25, the original frame was independently aligned to the robust global template, and the alignment yielding the higher correlation was retained. Bilinear interpolation with reflection padding was used to evaluate registration quality, whereas the iteratively corrected image stack was updated using nearest-neighbor interpolation with reflection padding.

**Multi-scale feature encoder** Following translation correction, dense non-rigid motion was estimated using PyLoReg-Net. The network received a reference template and a moving frame, each with dimensions (B×1×H×W). Both images were processed by a shared convolutional encoder consisting of an initial (3×3) convolution followed by four hierarchical downsampling stages. The initial convolution generated 64-channel features at the original image resolution. At each subsequent level, feature maps were downsampled by a factor of two using (2×2) average pooling and processed by a double-convolution block. Each block contained two (3×3) convolutions with reflection padding, instance normalization with learnable affine parameters, and ReLU activation. The downsampled feature levels contained 128 channels. The resulting feature pyramid therefore contained representations at the original, (1/2), (1/4), (1/8), and (1/16) spatial resolutions. For flow estimation, the feature pyramid was ordered from coarse to fine, and four scales were used by default: (1/16), (1/8), (1/4), and (1/2) of the original image resolution. An auxiliary decoder reconstructed each input image from the lowest-resolution feature map using four rounds of bilinear upsampling, (1×1) channel projection, and double-convolution refinement. The decoder contained no skip connections.

**Local correlation cost volumes** At each pyramid level, correspondence between the reference and moving feature maps was represented using a PWC-style local cost volume. Feature vectors were first normalized to unit _2_ norm along the channel dimension. For each spatial position in the reference feature map, similarity was calculated between the reference feature vector and moving-image feature vectors within a local neighborhood of radius four feature pixels. Similarity was computed as the channel-wise dot product between normalized feature vectors and was scaled by the square root of the feature dimension. This procedure generated a (9×9) search region containing 81 candidate displacements at each spatial location. Replicate padding was applied to the moving feature map before local neighborhoods were extracted using an unfolding operation. The resulting cost volume had dimensions (B×81×H_s_×W_s_), where (B) is the batch size and (H_s_) and (W_s_) are the height and width of the feature map at pyramid level (s). Because the search radius was fixed in feature-map coordinates, the corresponding search range in the original image decreased from coarse to fine levels. A radius of four feature pixels represented maximum local displacements of approximately 64, 32, 16, and 8 pixels at the (1/16), (1/8), (1/4), and (1/2) resolution levels, respectively.

**Coarse-to-fine flow estimation** At each scale, the 81-channel cost volume was passed to a convolutional refinement network to estimate a two-channel residual flow field. The refinement network contained three (3×3) convolutional layers with 128 output channels and ReLU activation, followed by a final (3×3) convolution producing the horizontal and vertical flow components. At the coarsest scale, the refinement network predicted an initial displacement field directly from the cost volume. At each subsequent scale, the cumulative flow estimate from the preceding coarser level was resized to the spatial dimensions of the current feature map using bilinear interpolation. The horizontal and vertical displacement components were rescaled according to the corresponding changes in feature-map width and height. The moving feature map was then warped using the resized cumulative flow, and a new local cost volume was computed between the reference feature map and the warped moving feature map. The refinement network predicted an additional residual flow field from this cost volume. This residual estimate was added to the upsampled cumulative flow, progressively refining the displacement field from coarse to fine spatial resolution.

**Learned flow upsampling** Flow fields were upsampled using a learned transposed-convolution module. The module supported spatial scale factors of 2, 4, 8, and 16. Its first layer mapped the two-channel flow field to 32 channels using a (4×4) transposed convolution with stride two and padding one, followed by LeakyReLU activation with a negative slope of 0.1. Additional layers retained 32 channels and applied the same twofold upsampling operation. A final (3×3) convolution projected the upsampled representation back to two flow channels. After spatial upsampling by a factor, the displacement amplitudes were multiplied by the factor so that the predicted flow remained expressed in pixels of the higher-resolution coordinate system. Residual flow predictions from each scale were also independently upsampled to full resolution and retained as intermediate outputs. All results reported here used transposed-convolution upsampling.

**Iterative registration and one-shot raw-data warping** After the initial rigid correction, two rounds of PyLoReg-Net-based dense registration were performed. At each round, a robust global template was regenerated from the currently corrected stack, and frames were processed in batches of two. Inference was performed in evaluation mode with automatic mixed precision when a CUDA-capable GPU was available. For quality control, motion magnitude was calculated from the horizontal and vertical displacement components, and frame-to-template similarity was measured by normalized cross-correlation on a centered 256 × 256-pixel crop sampled with stride two. The processed image stack was updated after each round using nearest-neighbor interpolation and reflection padding. Rigid and dense displacement fields from all rounds were composed into a single cumulative flow field using bilinear interpolation with zero padding. To reduce memory usage, cumulative flow components were stored as unsigned 16-bit integers using a scale factor of 100 and an offset of 32,767.5. The original raw stack was retained unchanged during iterative registration and warped only once using the final composed flow. The registered stack was then rescaled to the original intensity range, converted to unsigned 16-bit format, and saved as a TIFF file.

### Synthetic and experimental training data for PyLoReg-Net

To train and evaluate the motion correction model (PyLoReg-Net), we used a combination of large-scale synthetic datasets and experimentally acquired two-photon calcium imaging recordings.

**Synthetic data generation using NAOMi** Synthetic training data were generated using the NAOMi simulation framework^79^, which provides a physics-based pipeline for realistic two-photon microscopy data synthesis. The simulation pipeline consisted of five sequential modules: (1) Neural volume generation. Three-dimensional neural volumes were generated, including neuronal somata, dendrites, axons, vasculature, and background structures, with realistic spatial distributions. (2) Neural activity simulation. Temporal activity traces were generated using stochastic spiking models, producing neuron-specific calcium dynamics with configurable firing rates and temporal resolution. (3) Optical propagation modeling. Optical effects were simulated using point spread function (PSF) modeling based on system parameters such as numerical aperture (NA), excitation power, and imaging depth. (4) Scanning simulation. The neural volume and activity were combined through a simulated raster scanning process to generate time-lapse image sequences mimicking two-photon imaging acquisition. (5) Noise modeling. Realistic noise sources were incorporated during scanning, including photon shot noise and background fluctuations, to match the statistical properties of experimental data.

Using this pipeline, we generated large-scale synthetic datasets with varying imaging depths, neuronal densities, firing rates, and optical conditions. In our implementation, volumes of size up to 500 × 500 × 60 voxels were simulated, with temporal sequences of several thousand frames per recording. Multiple simulation runs were performed with varying parameters to increase diversity and improve model generalization.

**Experimental datasets** In addition to synthetic data, we incorporated multiple publicly available two-photon calcium imaging datasets for training and validation. These datasets span diverse brain regions, imaging conditions, and signal-to-noise regimes. A summary of the datasets used is provided in FigureS1D, including Neurofinder benchmark datasets and CRCNS two-photon datasets (pvc and ssc series). These datasets collectively comprise hundreds of imaging stacks and over one million frames, covering a wide range of imaging conditions.

### Neuron segmentation and trace extraction

Neuron segmentation and activity extraction were performed on NeuroPilot-processed image stacks using a neurite-aware pipeline based on Suite2p^26,27^. For each recording, maximum intensity projections (MIP) and standard deviation (STD) projections were computed to capture complementary structural and activity-related features, and segmentation was conducted with parameters optimized for elongated and overlapping structures. NeuroPilot correction and baseline normalization were applied to extract fluorescence activity traces. To improve segmentation quality, an intensity-based filtering step was applied using normalized MIP and STD images. For each ROI, mean intensity within its spatial footprint was computed, and ROIs that were simultaneously dim in both representations were removed, preserving structures that are either structurally prominent or functionally active. For downstream analysis, ROI masks were further refined by retaining the largest connected component and ranking ROIs based on the maximum amplitude of their fluorescence traces. ROIs were retained if they passed predefined intensity, size and connected-component criteria. No manual ROI addition was performed. When morphological refinement was applied, the same deterministic operation was applied to all ROIs in the corresponding dataset and recorded in the processing log. The resulting ROI masks and ΔF/F traces were used for all subsequent analyses.

### Cross-session and longitudinal alignment

To enable consistent tracking of neural activity across sessions and over extended timescales, we performed cross-session alignment using a two-stage pipeline combining image restoration and optical flow–based registration. For each imaging session, raw image stacks were first processed using NeuroPilot to remove noise and correct intra-session motion artifacts, yielding denoised and motion-stabilized recordings. From each processed stack, the STD projection and MIP were computed, capturing complementary functional and structural features, respectively.

Dense non-rigid transformations between sessions were estimated using PyLoReg-Net applied to pairs of MIPs. These projections provide robust representations for cross-session correspondence, allowing accurate estimation of spatial deformation fields even under substantial variability in signal intensity and activity patterns. The resulting optical flow fields encode pixel-wise displacements between sessions. The estimated deformation fields were subsequently applied to the full spatiotemporal image stacks by warping each frame to a common reference coordinate system. This strategy enables efficient and consistent alignment of entire recordings while preserving temporal dynamics. The resulting dataset is fully aligned across sessions, enabling direct comparison of neuronal activity over time and supporting downstream analyses such as longitudinal neuron tracking and population-level dynamics.

### Imaging datasets and experimental conditions

**One-photon miniscope imaging** We analyzed a previously published MiniXL dataset acquired from the dorsal hippocampal CA1 region during linear-track behavior. GCaMP6f-expressing neurons in the dCA1 pyramidal layer were imaged through a stacked pair of 1.8-mm-diameter GRIN lenses using a MiniXL one-photon epifluorescence miniscope. The lower 4.7-mm-long GRIN lens was implanted above dCA1, with its center positioned at AP −2.1 mm, ML 1.8 mm, and DV −1.25 mm relative to bregma. Calcium imaging was performed at 23 frames/s for 15 min per session while mice traversed a 2-m linear track. Three sessions were acquired at 2-day intervals. Raw recordings were collected at 1,000 × 1,000 pixels and cropped to 704 × 704 pixels for analysis, corresponding to an approximately 1.393 × 1.393 mm FOV. Behavioral videos were recorded simultaneously at 50 frames/s.

**Sciatic nerve injury model** To induce neuropathic pain, the sciatic nerve injury (SNI) model was established as previously described. Mice were first subjected to baseline imaging (defined as Day 1 in this study). Immediately following imaging, animals underwent SNI surgery. Briefly, mice were deeply anesthetized with 1.5–3% isoflurane delivered in 100% oxygen. After sterilization with 70% ethanol, animals were placed in a prone position and the limbs were secured to an anatomical plate. A dorsal longitudinal skin incision was made spanning vertebral levels T11 to L1 to expose the underlying musculature. The biceps femoris muscle was bluntly separated using forceps to reveal the sciatic nerve and its three terminal branches: the sural, common peroneal, and tibial nerves. Neuropathic injury was induced by tightly ligating the common peroneal and tibial nerves using 5-0 silk sutures (RockTech), followed by transection distal to the ligation site and removal of approximately 4 mm of the distal nerve stump. The sural nerve was left intact. The incision site was then closed and sutured using 5-0 silk sutures.

### Behavioral decoding and decoder-derived similarity analysis

To characterize the association between neural population activity and behavioral states, we combined supervised behavioral decoding with decoder-derived similarity analysis leveraging confusion structure and population statistics.

**Data preprocessing and feature construction** Neuronal activity traces were converted to ΔF/F traces and structured as a time-by-neuron matrix. To maintain computational tractability and ensure balanced sampling across extended recordings, the data were temporally subsampled to a fixed resolution. Each time point was associated with a behavioral label derived from synchronized annotations, with categorical states encoded as integer indices. Prior to decoding, neural features were z-score normalized using statistics derived exclusively from the training set; these identical parameters were then applied to the test set to preclude data leakage.

**Train-test splitting with temporal structure preservation** To evaluate decoding performance while accounting for temporal dependencies, we implemented a block-wise train-test splitting strategy. The time series was partitioned into contiguous blocks, a subset of which was allocated to the test set. To ensure evaluative robustness, we required all behavioral classes to be represented in both partitions. For analyses requiring stratified random splitting, decoding results were treated as exploratory and were not used as primary evidence for state separability.

**Behavioral decoding models** Behavioral states were decoded from neural population activity using a linear support vector machine (LinearSVM). The classifier was trained to predict behavioral labels based on instantaneous population activity vectors. A linear decoder was selected to provide an interpretable measure of population-level separability while mitigating the risk of overfitting in high-dimensional neural data. Class imbalance was addressed via class-weighting where appropriate. Decoding performance was assessed on held-out test data using multiple metrics, including overall accuracy, balanced accuracy, and the macro-averaged F1 score.

**Decoder-derived behavioral similarity analysis** To characterize the structure of decoding errors and identify relationships between behavioral states, we analyzed confusion matrices derived from model predictions. The matrices were normalized on a per-class basis to estimate the conditional probability of predicted states given the ground truth. These normalized matrices capture the representational similarity between neural population activity patterns across different behaviors. To formally quantify this similarity, we constructed a symmetric similarity matrix:

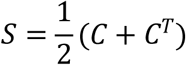

where denotes the normalized confusion matrix. Hierarchical clustering was then applied to this similarity matrix to identify groups of behaviors with similar neural representations.

**Directed confusion structure and asymmetry analysis** To further characterize directional relationships between behavioral representations, we computed an asymmetry matrix:

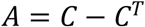

which quantifies directional biases in misclassification between behavioral states. We additionally constructed directed confusion graphs, where nodes represent behavioral states and edges correspond to dominant misclassification pathways. Edge weights were derived from the largest off-diagonal elements of the confusion matrix. Chord diagrams were used to visualize these directed relationships in a compact form.

### Behavioral state-transition analysis

To characterize neural population dynamics associated with transitions between behavioral states, frame-wise behavioral annotations were first divided into contiguous episodes with identical state labels. A behavioral transition was defined as the boundary between two consecutive episodes assigned to different states, and transition onset was designated as the first frame of the target-state episode. We analyzed six prespecified transitions: NREM-to-REM, REM-to-NREM, nest grooming-to-NREM, NREM-to-nest grooming, motor-to-locomotion, and locomotion-to-motor.

For each transition event, neuronal activity was extracted within a 120-s window extending from 60 s before to 60 s after transition onset. Time zero corresponded to the first frame labeled as the target behavioral state. Events were included only when the complete analysis window was contained within the recording and the contiguous source and target episodes each lasted for at least 5 s. Transition classes containing fewer than three valid events were excluded from subsequent analyses.

Neuronal fluorescence traces were converted to ΔF/F and organized as a time-by-neuron matrix. For transition-aligned analyses, each neuron’s activity was standardized across the complete recording by subtracting its temporal mean and dividing by its temporal standard deviation. Event-aligned activity matrices were retained at the native temporal resolution, without temporal interpolation.

To define representative population activity patterns for individual behavioral states, stable-state frames were identified as frames located at least 30 s from any annotated behavioral transition. For each state represented by at least 50 stable frames, a population centroid was calculated as the mean standardized activity vector across all corresponding stable frames. At each transition-relative time point, Euclidean distances from the instantaneous population activity vector to the source-and target-state centroids were calculated. Distance trajectories were first calculated separately for individual transition events and were subsequently summarized across events.

Population trajectories around each transition were additionally examined in principal-component space. A PCA model was fitted using stable-state population activity, with up to six principal components retained, and the learned transformation was then applied to the complete recording. PCA trajectories were aligned to transition onset and averaged across events belonging to the same transition class. The first two principal components were used to visualize the progression of neural population activity from the source-state representation toward the target-state representation.

Transition-aligned mean population activity was calculated by averaging standardized activity across neurons at each time point. Population variability was quantified from the dispersion of neural activity around the transition. For visualization, time-resolved curves were smoothed using a one-dimensional Gaussian filter with a standard deviation of 3 s. Unsmoothed activity values were retained for numerical analyses.

To quantify changes in coordinated population activity, pairwise neuronal correlations were calculated within a 30-s temporal window centered on each transition-relative time point. A neuron-by-neuron Pearson correlation matrix was calculated for each event and temporal window, and the mean of the off-diagonal correlation coefficients was used as the population-wide correlation measure. Correlation time courses were subsequently averaged across events of the same transition type.

As a control for spontaneous fluctuations within the source behavioral state, source-state-matched pseudo-transition events were generated. For each genuine transition, a pseudo-transition center was randomly selected from frames assigned to the same source state. Candidate pseudo-transition centers were required to accommodate the complete 120-s analysis window and to occur at least 60 s from any genuine behavioral transition. Neural activity, PCA trajectories, and distances to source- and target-state centroids were calculated for pseudo-transition events using the same procedures as for genuine transitions.

The robustness of transition-aligned dynamics to uncertainty in behavioral boundary assignment was further assessed by temporally shifting transition centers by −60, −30, −15, +15, +30, or +60 s and repeating the alignment procedure. Unless otherwise indicated, transition-aligned measurements are presented as the mean ± SEM across individual transition events, with each transition event treated as the unit of analysis.

### Dynamical systems modeling based on rSLDS

To characterize the temporal organization and dynamical structure of neuronal population activity, we employed a recurrent switching linear dynamical system (rSLDS), which approximates nonlinear dynamics as a latent mixture of locally linear regimes. Neuronal activity traces (ΔF/F) were first projected into a lower-dimensional observation space using Principal Component Analysis (PCA). Specifically, the population activity matrix (time × neurons) was decomposed into its top principal components (typically 10), preserving dominant variance while attenuating noise and redundancy. These reduced observations were then modeled via the *ssm* framework. Within this rSLDS architecture, neural population trajectories transition through a set of discrete dynamical states, each governed by an underlying linear dynamical system. At each time point, the latent state ∈ evolves according to:

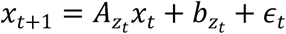

Where *Z_t_* ∈ {1, …, *K*} denotes the discrete state, *A_zt_* and *b_zt_* define the linear dynamics for that state, and ∈_t_ is Gaussian noise. The discrete state transitions are governed by a recurrent transition model that depends on the continuous latent state, enabling context-dependent switching between dynamical regimes. Model fitting was performed using variational inference (Laplace-EM), with latent dimensionality set to 2 for visualization and interpretation of population trajectories. Initialization was performed using PCA-based projections to ensure stable convergence.

After training, we extracted both the continuous latent trajectories x_t_ and the most likely discrete state sequence *z*_t_. The learned system matrices *A*_k_ and offsets for each discrete state were directly retrieved from the trained model. To visualize the underlying dynamics, we computed the velocity field associated with each dynamical regime. Specifically, the local flow at state x was approximated as:

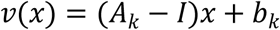

which represents the instantaneous displacement of the latent state under the learned dynamics. Velocity fields were evaluated over a grid spanning the latent space and visualized using vector field plots. Latent trajectories were overlaid onto these fields, allowing direct comparison between observed neural dynamics and the inferred dynamical structure. Trajectories were further color-coded either by behavioral state or by normalized time to reveal state-dependent and temporal evolution patterns.

To summarize global dynamics, we additionally computed an occupancy-weighted mixture flow field, in which the velocity fields of individual modes were combined according to their empirical usage frequencies:

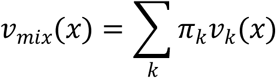

Where π_k_ denotes the fraction of time spent in each discrete state.

## Supporting information

NeuroPilot Stabilization of Spinal Cord Calcium Imaging across Stimulation and Behavioral Conditions

Longitudinal NeuroPilot Tracking of Spinal Cord across 28 Days in an SNI Model

Continuous 24h mPFC Calcium Imaging Aligned with Spontaneous Behavior after NeuroPilot Processing

mPFC Population Activity and Latent Dynamics during NREM-to-REM Sleep Transitions

## Data and code availability

The NeuroPilot source code is publicly available on GitHub at https://github.com/LuoYuhong123/NeuroPilot and archived on Zenodo at https://zenodo.org/records/21835435. Analysis scripts, preprocessed datasets, and the interactive web implementation are available at https://neuropilot.it.com/. Additional raw imaging data supporting this study are available from the Lead Contact upon reasonable request, subject to applicable data-sharing and institutional requirements. The manuscript and supplementary materials provide the information required to reproduce the principal analyses.

## Acknowledgments

This work was supported by the Fundamental and Interdisciplinary Disciplines Breakthrough Plan of the Ministry of Education of China (JYB2025XDXM504), the National Natural Science Foundation of China (62401156), and the General Research Fund of the Research Grants Council of Hong Kong (RGC/GRF; project nos. 14119125, 14119024, and 14117122). We thank Beijing Transcend Vivoscope Bio-Technology Co., Ltd., Nanjing Raygen Health Technology Co., Ltd., and TRANSVISTA for insightful technical discussions that contributed to the development of this work.

## Author contributions

Z.L., N.C., X.L., and G.Z. supervised the project. G.Z. and Y.L. developed the conceptual and technical architecture of NeuroPilot, established the overall computational workflow, and implemented its core components. G.Z. and Z.L. led the data analysis and interpretation of the results and prepared all main and supplementary figures. Y.L. and J.W. produced the supplementary videos. Y.L. and D.L. developed and deployed the web-based implementation, with web design support from T.W. F.J. and P.W. advised on the analysis of head-mounted imaging data and contributed the spinal cord and deep-brain imaging datasets. D.Z. and J.H. contributed the three-photon imaging datasets and provided scientific and technical feedback. S.Q., C.G., and B.L. contributed the mitochondrial calcium, one-photon, and large-FOV two-photon imaging datasets, respectively. M.M. advised on the attractor analysis. X.L. and Y.J. provided guidance on the analysis and interpretation of the spinal cord imaging data. G.Z., Z.L., and X.L. led the preparation of the manuscript. All authors discussed the results, reviewed the manuscript, and approved the final version.

