## Supplementary Information for "NeuroPilot: Longitudinal intravital neural imaging across scales and modalities through an agentic platform"

#### 4 Supplementary Videos

| | |
|---|---|
| **Video S1** | NeuroPilot enables context-aware processing and analysis of large-scale neural imaging data, related to Figure 1.<br>This video presents the NeuroPilot workflow for processing heterogeneous and longitudinal neural imaging datasets. NeuroPilot coordinates denoising, non-rigid motion correction, neuronal tracking, and data interpretation through a context-aware workflow. The video also introduces PyLoReg-Net, a coarse-to-fine optical-flow module for pixel-wise motion correction. |
| **Video S2** | NeuroPilot stabilization of spinal cord calcium imaging across behavioral and sensory conditions, related to Figure 3.<br>Representative spinal cord calcium imaging recordings acquired under spinal fixation, freely moving behavior, cold stimulation, nociceptive mechanical stimulation by hind-paw pinch, and air-puff stimulation. For each condition, the raw recording and the corresponding NeuroPilot-processed recording are shown together with synchronized kymographs. Videos are displayed at 30× real time. Scale bars, 50 μm for the full FOV and 10 μm for the magnified regions. |
| **Video S3** | Longitudinal NeuroPilot tracking of spinal cord neurons across 28 days in an SNI model, related to Figure 3 and Figure S5.<br>Representative miniature two-photon calcium imaging recordings from the L3–L5 spinal cord segments of freely moving mice are shown across five longitudinal sessions: Day 1 before injury and Days 5, 14, 21, and 28 after SNI. Raw and NeuroPilot-processed recordings are displayed for comparison. In the first portion, imaging sequences are displayed at 20× real time. Scale bars, 50 μm for the full FOV and 25 μm for the magnified regions. The later portion shows STD projection images overlaid with color-coded neuronal footprints, followed by synchronized calcium traces from representative neurons matched across all five sessions. Traces extracted from raw recordings are shown in white, and those extracted from NeuroPilot-processed recordings are shown in green. The later portion is displayed at 30× real time. |
| **Video S4** | NeuroPilot restoration across deep-brain, large-field, and volumetric calcium imaging, related to Figure 4, Figure S8, and Figure S7.<br>Miniature two-photon calcium imaging through GRIN lenses is shown for two deep-brain regions, the lateral entorhinal cortex (LEC; DV = 4.30 mm) and hippocampal CA1 (DV = 1.20 mm). Raw and NeuroPilot-processed |

| | |
|---|---|
| | recordings are displayed side by side, highlighting improved neuronal visibility and temporal signal recovery. Videos are shown at 12× real time. Scale bars, 25 μm. Large-field cortical two-photon imaging is then presented across an approximately 6-mm FOV, together with magnified views of three local regions. Raw and NeuroPilot-processed recordings are shown for comparison. Videos are displayed at 10× real time. Scale bar, 1 mm. The video concludes with volumetric head-mounted three-photon calcium imaging extending through the cortex and into hippocampal CA1. Three-dimensional renderings of the raw and NeuroPilot-processed volumes are shown side by side, illustrating restoration of low-SNR signals and improved neuronal visibility across imaging depth. |
| **Video S5** | NeuroPilot restoration of multiplexed, vascular, and mitochondrial imaging data, related to Figure S7.<br>Dual-channel two-photon imaging of the mouse vibrissal primary somatosensory cortex (vS1) is shown with GCaMP6s calcium signals in green and mCherry-labeled structural signals in magenta. Raw and NeuroPilot-processed recordings are displayed side by side, with magnified regions highlighting recovery of neuronal structure and temporal signal definition in both channels. Videos are displayed at 20× real time. Scale bars, 50 μm for the full FOV and 25 μm for the magnified regions. Spinal cord vascular imaging from freely moving mice is then shown, comparing raw and NeuroPilot-processed recordings and highlighting improved continuity and stability of the microvascular network. Videos are displayed at 9× real time. Scale bars, 50 μm for the full FOV and 10 μm for the magnified regions. The video concludes with two-photon imaging of mitochondrial calcium dynamics and network morphology. Raw and NeuroPilot-processed fluorescence recordings are shown together with magnified regions illustrating recovery of continuous tubular structures, followed by skeletonization analysis of the reconstructed mitochondrial network. Videos are displayed at 19.5× real time. Scale bars, 50 μm for the full FOV and 25 μm for the magnified regions. |
| **Video S6** | Continuous 24-hour mPFC calcium imaging aligned with spontaneous behavior after NeuroPilot processing, related to Figure 5 and Figure S9.<br>Continuous head-mounted two-photon calcium imaging of the medial prefrontal cortex (mPFC) in a freely behaving mouse is shown across 24 hours. NeuroPilot-processed imaging is displayed together with synchronized infrared behavioral recordings, segmented neuronal |

| | |
|---|---|
| | footprints, population calcium activity heatmaps, and a circular behavioral ethogram. Color-coded annotations indicate spontaneous behavioral and sleep states throughout the recording. Videos are displayed at 120× real time. Scale bars, 50 μm for the full FOV and 20 μm for the magnified regions. |
| **Video S7** | mPFC population activity and latent dynamics during spontaneous NREM-to-REM sleep transitions, related to Figure 6 and Figure S10.<br>Representative spontaneous NREM-to-REM transitions are shown from continuous head-mounted two-photon calcium imaging of the mPFC in freely behaving mice. NeuroPilot-processed imaging is displayed together with synchronized behavioral-state annotations, segmented neuronal ROIs, z-scored population calcium activity, and rSLDS-inferred latent trajectories. The aligned views illustrate the reorganization of mPFC population activity as neural dynamics evolve from NREM sleep into REM sleep. Videos are displayed at 60× real time. Scale bar, 50 μm. |

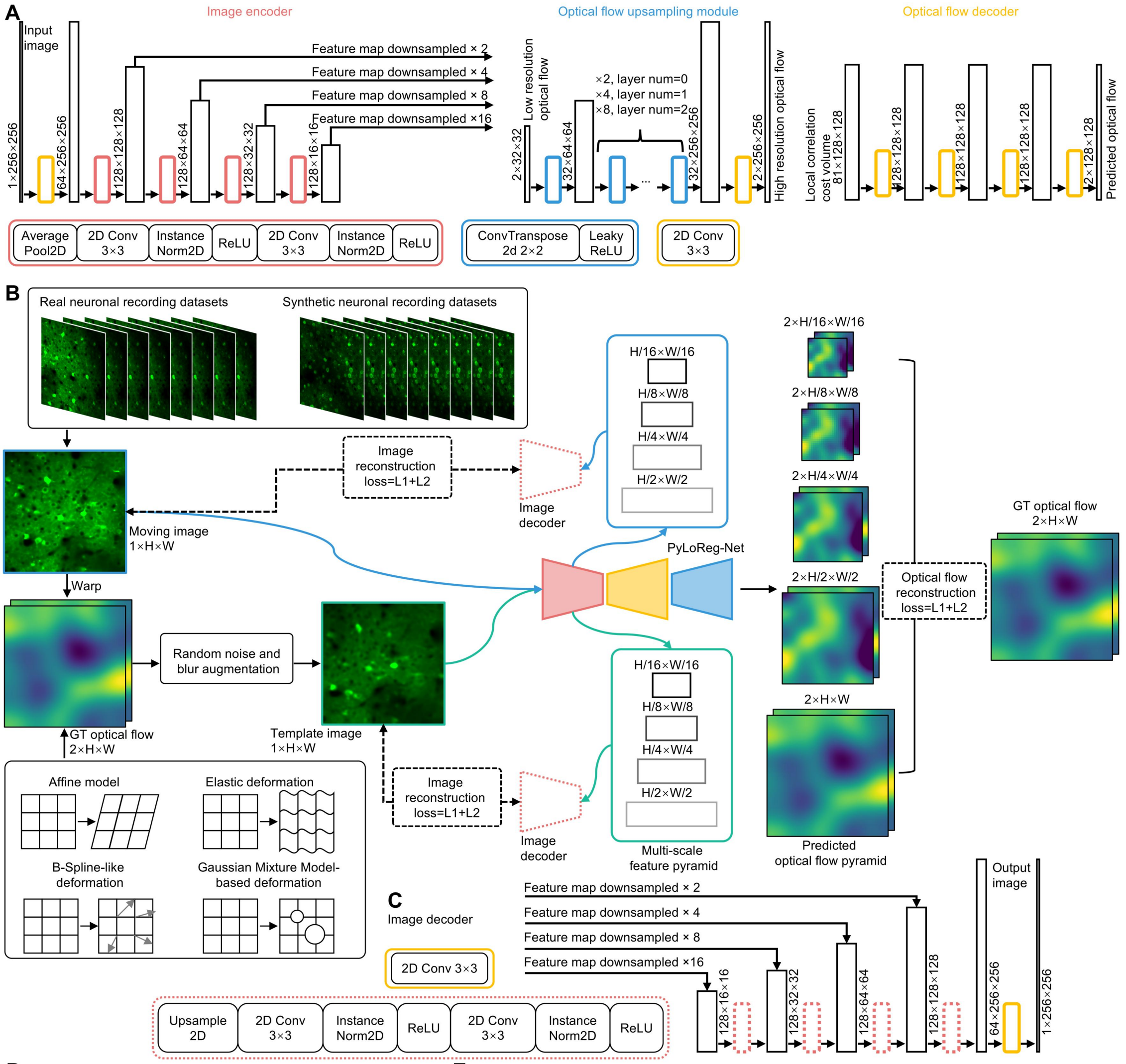


D

| Dataset name | Number of stacks | Number of frames | Image size/pixel | Usage in training | Notes |
|---|---|---|---|---|---|
| Synthetic dataset | 200 | 200,000 | 490×490 | All for training | 90.5 GB |
| Neuronfinder | 28 | 103,517 | 512×512 | All for training | 100.7 GB |
| CRCNS pvc-13 | 44 | 97,511 | 256×512 | All for training | 22.7 GB |
| CRCNS pvc-10 | 116 | ~541,000 | 256×455 | Part for training | 234 GB |
| CRCNS ssc-8 | 53 | 107,417 | 128×256 | All for training | 6.9 GB |
| CRCNS ssc-1 | 125 | ~350,000 | 512×512 | Part for training | 341 GB |
| CRCNS ssc-2 | 258 | ~558,000 | 512×512 | Part for training | 543 GB |
| CRCNS ssc-6 | 10 | 166,600 | 128×512 | All for training | 10 GB |

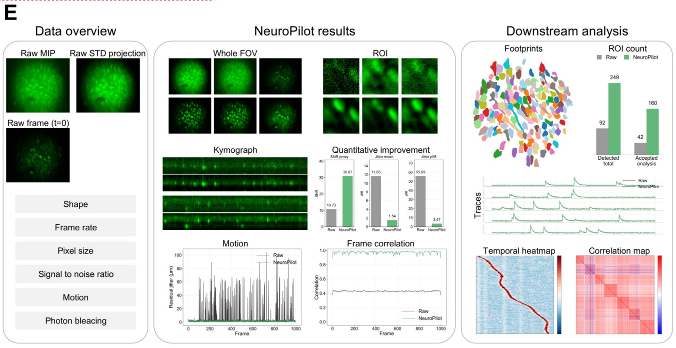


**Figure S1**

**PyLoReg-Net architecture, training strategy, and datasets for non-rigid motion correction in NeuroPilot.**

(A) PyLoReg-Net architecture for dense non-rigid motion estimation. The encoder extracts multi-scale image features at 2×, 4×, 8×, and 16× downsampling. These features are used to estimate low-resolution optical flow, which is progressively upsampled and refined to produce a dense displacement field.

(B) Training framework for PyLoReg-Net. The network is trained on experimentally acquired and synthetic calcium imaging recordings. Synthetic image pairs are generated with ground-truth (GT) displacement fields and noise/blur augmentation. Training combines flow reconstruction with auxiliary image reconstruction from decoded multi-scale features.

(C) Image decoder for the auxiliary reconstruction objective. Multi-scale encoder features are progressively upsampled and decoded into an image, which is compared with the corresponding target image using an L1 + L2 reconstruction loss.

(D) Calcium imaging datasets used for training. The training corpus includes a synthetic dataset generated with NAOMi and multiple experimentally acquired datasets from public repositories, including Neurofinder and CRCNS. The table summarizes dataset size, image dimensions, training usage, and data volume.

(E) Schematic overview of the traceable NeuroPilot report. The report integrates raw data summaries, restored imaging outputs, motion-correction metrics, segmented footprints, calcium traces, temporal heatmaps, and correlation maps for downstream analysis.

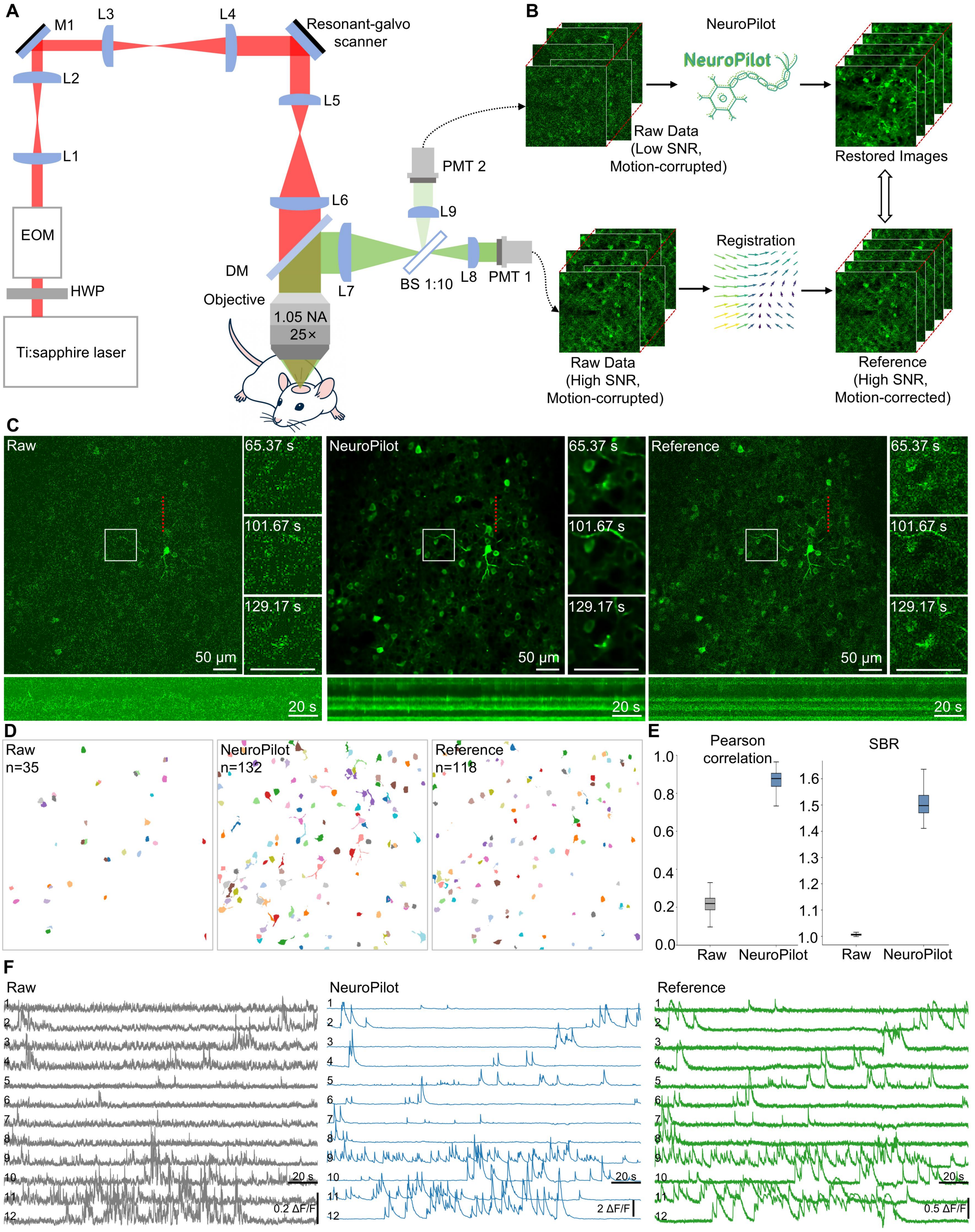


**Figure S2**

**Two-photon imaging setup, paired validation workflow, and quantitative assessment of NeuroPilot.**

(A) Schematic of the two-photon microscopy system used to acquire paired low- and high-SNR calcium imaging recordings. A Ti laser is modulated by a half-wave plate (HWP) and an electro-optic modulator (EOM), relayed through the scan optics, and delivered to the cortex through a resonant-galvo scanner and objective. Emitted fluorescence is collected through the objective, separated by a dichroic mirror (DM), split by a beamsplitter (BS), and detected by two photomultiplier tubes (PMTs) to generate paired low- and high-SNR recordings. L1–L9, lenses; M1, mirror.

(B) Paired validation workflow. Motion-corrupted high-SNR recordings are motion corrected to generate a high-SNR reference. The corresponding low-SNR recordings are processed by NeuroPilot, and the restored data are quantitatively compared with the high-SNR reference.

(C) Representative FOV comparing raw low-SNR recordings, NeuroPilot-restored data, and high-SNR reference recordings from in vivo imaging of neuronal somata. White boxes indicate regions shown at higher magnification in the adjacent time-lapse insets at the indicated time points. Spatiotemporal kymographs from the dashed red lines are shown below each field of view. Scale bars, 50 μm for FOV and 20 s for kymographs.

(D) Spatial footprints of neuronal somata segmented by Suite2p from raw low-SNR recordings, NeuroPilot-restored data, and high-SNR reference recordings. The number of segmented footprints is 35, 132, and 118 for raw, NeuroPilot-restored, and reference data, respectively.

(E) Quantitative comparison of raw and NeuroPilot-restored data against the high-SNR reference. Box plots show Pearson correlation coefficients between extracted calcium traces and reference traces, and SBRs of calcium traces extracted from raw and NeuroPilot-restored data.

(F) Representative calcium activity traces from 12 selected neuronal somata in raw low-SNR recordings, NeuroPilot-restored data, and high-SNR reference recordings. Activity is shown as $\Delta F/F$. Scale bars, 20 s; vertical scale bars, 0.2 $\Delta F/F$ for raw, 2 $\Delta F/F$ for NeuroPilot-restored data, and 0.5 $\Delta F/F$ for reference data.

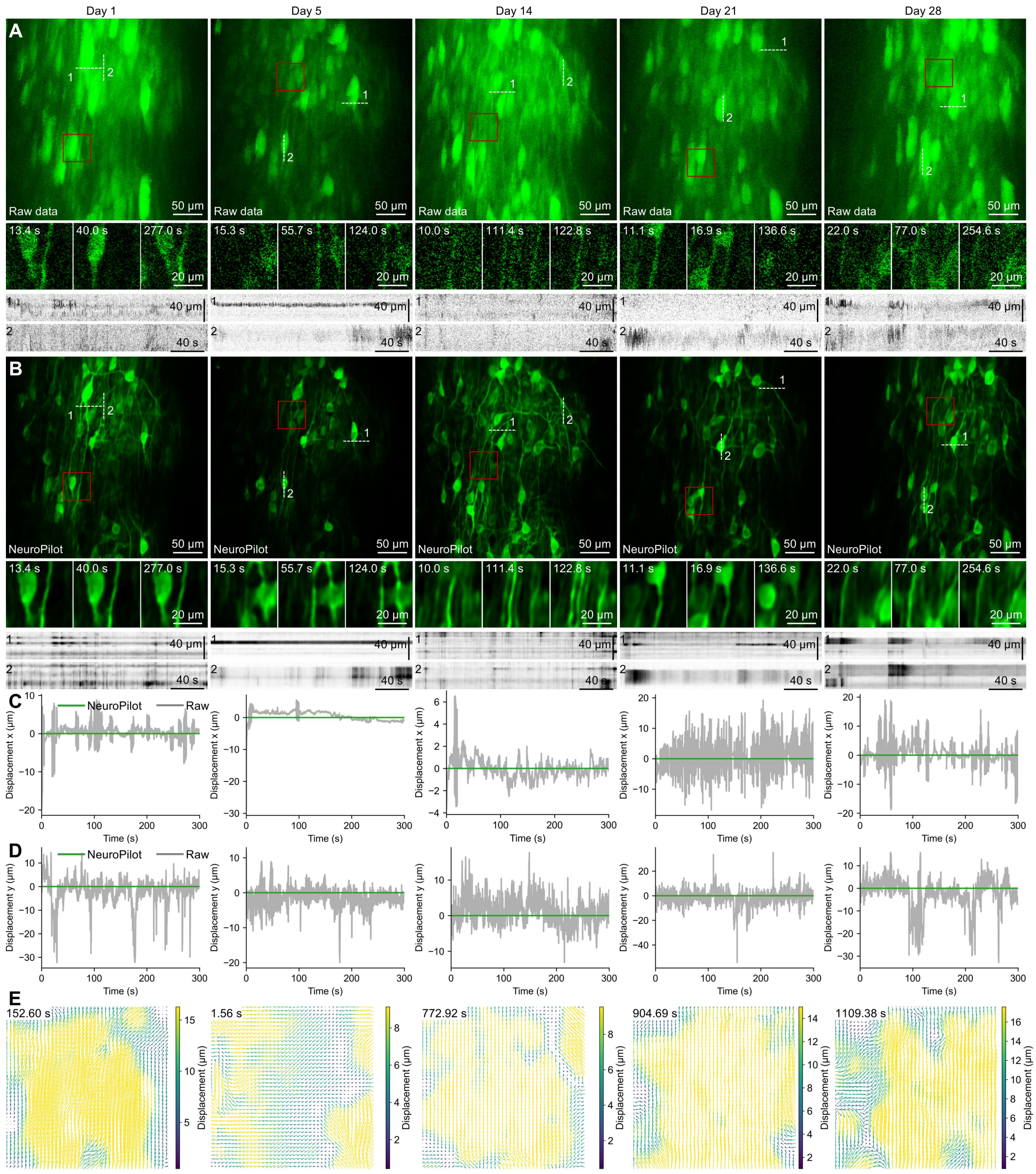


**Figure S3**

**Cross-day NeuroPilot stabilization of chronic spinal cord imaging in freely moving mice.**

(A-B) Representative longitudinal calcium imaging recordings from the L3-L5 spinal cord segments across five sessions in an SNI model. Day 1 corresponds to the pre-injury baseline, and days 5, 14, 21, and 28 correspond to post-SNI sessions. MIPs of raw data are shown in (A), and NeuroPilot-processed data are shown in (B). Bottom panels show magnified time-lapse frames from the red-boxed regions and spatiotemporal kymographs extracted along the dashed lines labeled 1 and 2. Scale bars, 50 μm for MIPs; 20 μm for insets; 40 μm and 40 s for kymographs.

(C-D) Estimated frame-by-frame displacements along the x axis (C) and y axis (D) before and after NeuroPilot processing. Gray, raw data; green, NeuroPilot-processed data. (E) Representative displacement fields estimated from individual sessions. Arrows indicate displacement direction, and color indicates displacement magnitude.

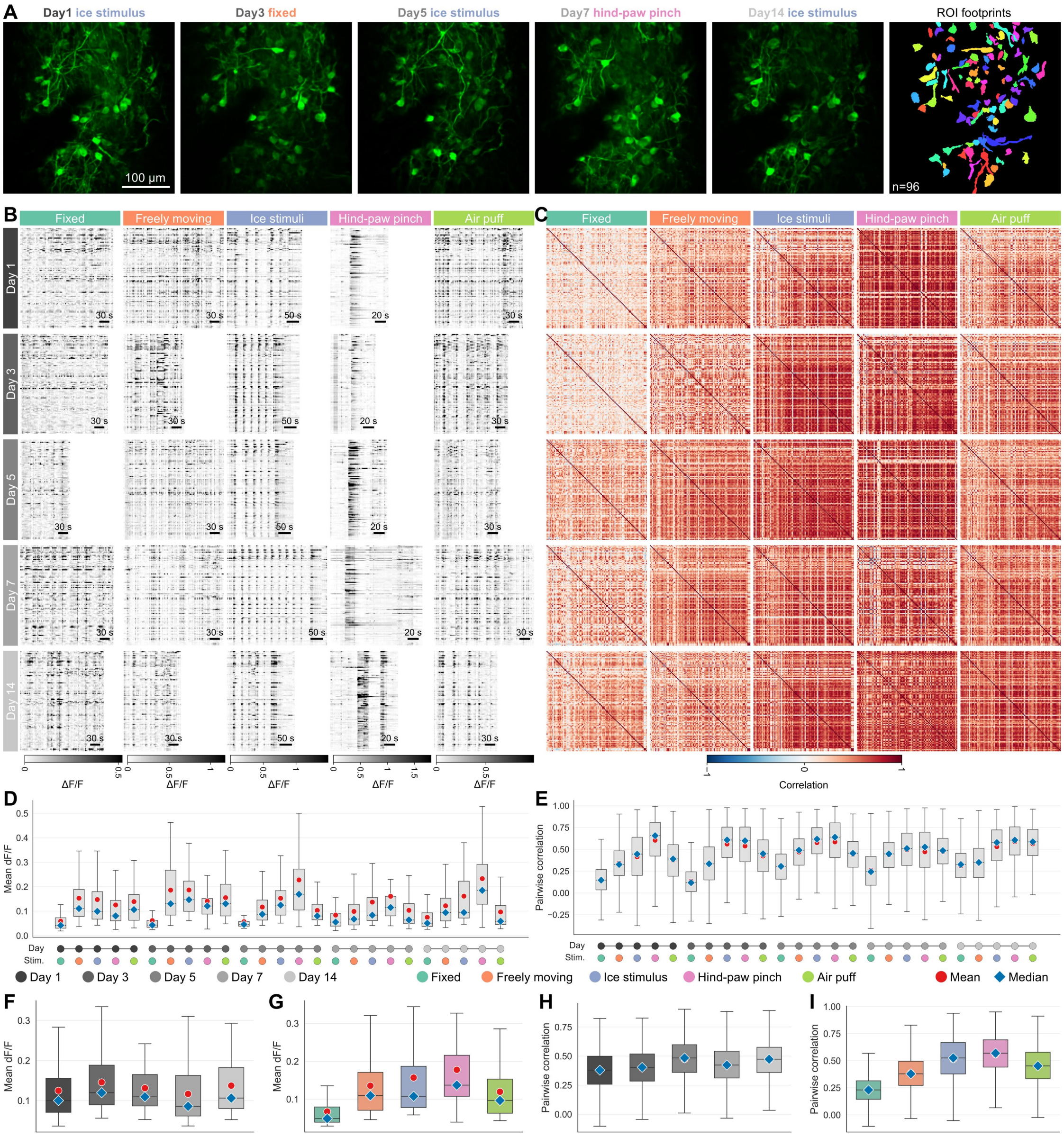


**Figure S4**

**Longitudinal tracking reveals stimulus-dependent spinal cord population responses across days.**

(A) Representative registered spinal cord FOV from the same animal across five imaging days and selected behavioral or sensory conditions. The rightmost panel shows the neuronal footprints used for longitudinal tracking. Scale bar, 100 µm; n = 96 tracked neurons.

(B) Activity heatmaps of longitudinally tracked ROIs across imaging days and experimental conditions. Within each heatmap, each row represents one tracked neuron. Conditions include spinal column-fixed, freely moving, cold stimulation, hind-paw pinch, and air puff. Activity is displayed as ΔF/F; within each condition column, all days are shown using the same ΔF/F color scale. Scale bars indicate time.

(C) Pairwise correlation matrices computed from the activity traces of tracked neurons for each day and condition shown in (B).

(D-E) Distributions of mean ΔF/F (D) and pairwise activity correlation (E) across each day-condition combination. Red circles indicate means, and blue diamonds indicate medians.

(F-G) Mean ΔF/F grouped by imaging day (F) or stimulus condition (G).

(H and I) Pairwise activity correlations grouped by imaging day (H) or stimulus condition (I).

Box plots show the median, interquartile range, and whiskers indicating the data range. Day and stimulus identities are indicated by the dot annotations below each plot. Fixed, spinal column-fixed condition; cold stimulation, ice-evoked cold stimulation.

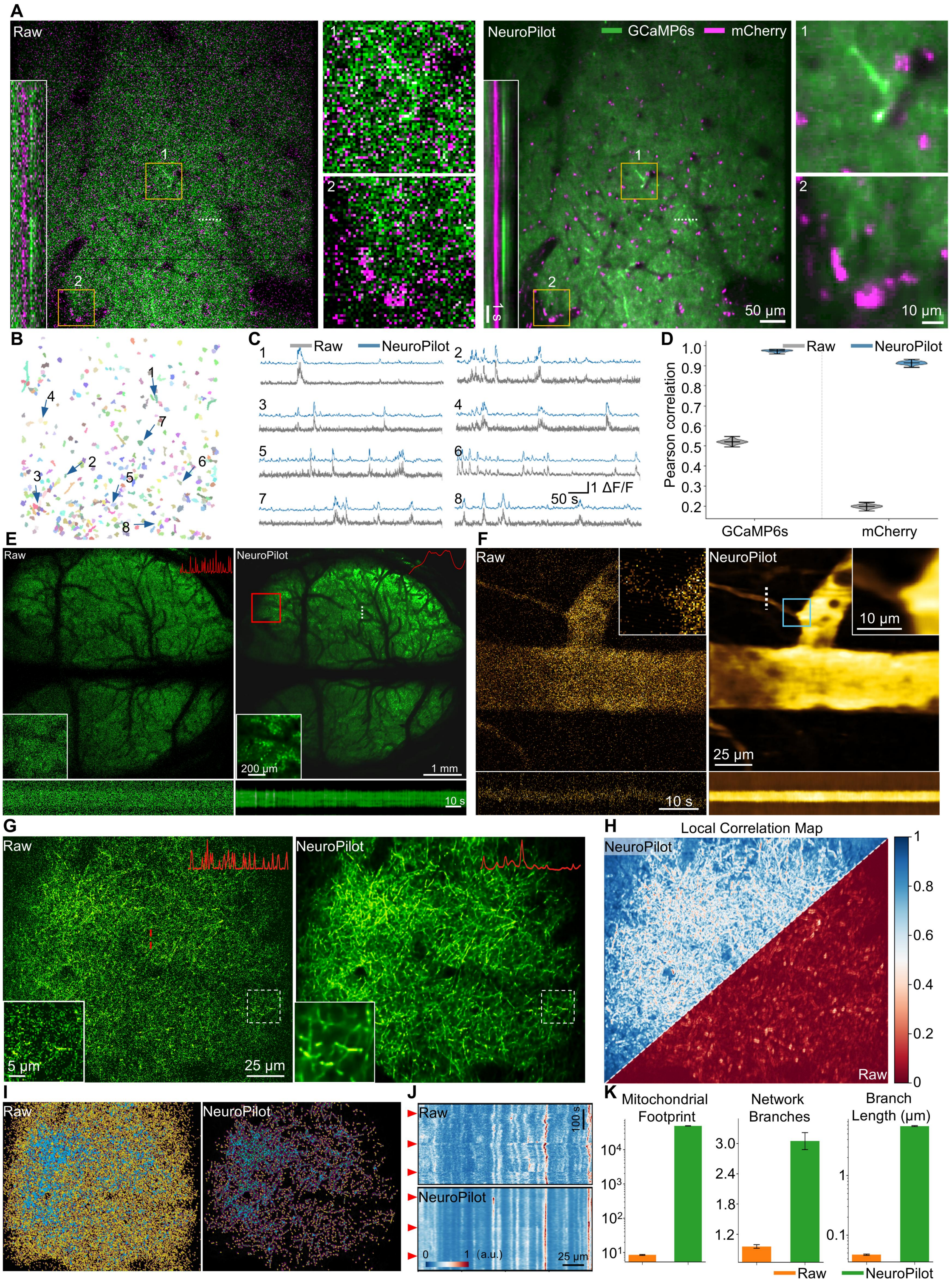


### Figure S5

**NeuroPilot restoration across multiplexed imaging channels and diverse biological targets.**

(A) Representative dual-channel two-photon imaging of mouse vibrissal primary somatosensory cortex (vS1) acquired at 7 Hz. Raw and NeuroPilot-restored images show

GCaMP6s signals in green and mCherry signals in magenta. Boxed regions indicate magnified insets. Kymographs extracted along the indicated dashed lines show temporal signal patterns before and after NeuroPilot restoration. Scale bars, 50 μm for main FOV, 10 μm for insets, and 1 s for kymographs.

(B) Spatial footprints of segmented neuronal ROIs across the FOV. Each pseudocolor denotes one segmented neuron.

(C) Representative GCaMP6s calcium fluorescence traces from selected neuronal ROIs. Traces correspond to the numbered ROIs in (B). Raw traces are shown in gray, and NeuroPilot-processed traces are shown in blue. Scale bars, 50 s and 1 $\Delta F/F$.

(D) Signal consistency analysis for the dual-channel dataset. Violin plots show Pearson correlation coefficients between raw and NeuroPilot-processed fluorescence traces for the GCaMP6s and mCherry channels.

(E) Representative large-field imaging example comparing raw and NeuroPilot-processed data. Insets show magnified regions, and kymographs extracted along the indicated dashed lines are shown below. Scale bars, 1 mm for the main FOV, 200 μm for the inset, and 10 s for the kymograph.

(F) Representative vascular imaging example comparing raw and NeuroPilot-processed data. Insets show magnified regions, and kymographs extracted along the indicated dashed lines are shown below. Scale bars, 25 μm for the main FOV, 10 μm for the inset, and 10 s for the kymograph.

(G) Representative mitochondrial imaging example comparing raw and NeuroPilot-processed data. Insets show magnified views of the dashed regions. Red traces indicate intensity profiles extracted along the indicated lines. Scale bars, 25 μm for the main FOV and 5 μm for the inset.

(H) Local correlation maps comparing raw and NeuroPilot-processed mitochondrial imaging data. The lower-right triangle shows raw data, and the upper-left triangle shows NeuroPilot-processed data.

(I) Mitochondrial segmentation and skeletonization using the MINA pipeline. Distinct colors denote segmented mitochondrial structures and skeleton features in raw and NeuroPilot-processed data.

(J) Spatiotemporal intensity profiles extracted along the indicated lines in mitochondrial recordings. Red arrowheads indicate time points with pronounced motion artifacts in the raw recordings. Scale bars, 100 s and 25 μm.

(K) Quantification of mitochondrial morphology from raw and NeuroPilot-processed data, including mitochondrial footprint, number of network branches, and branch length.

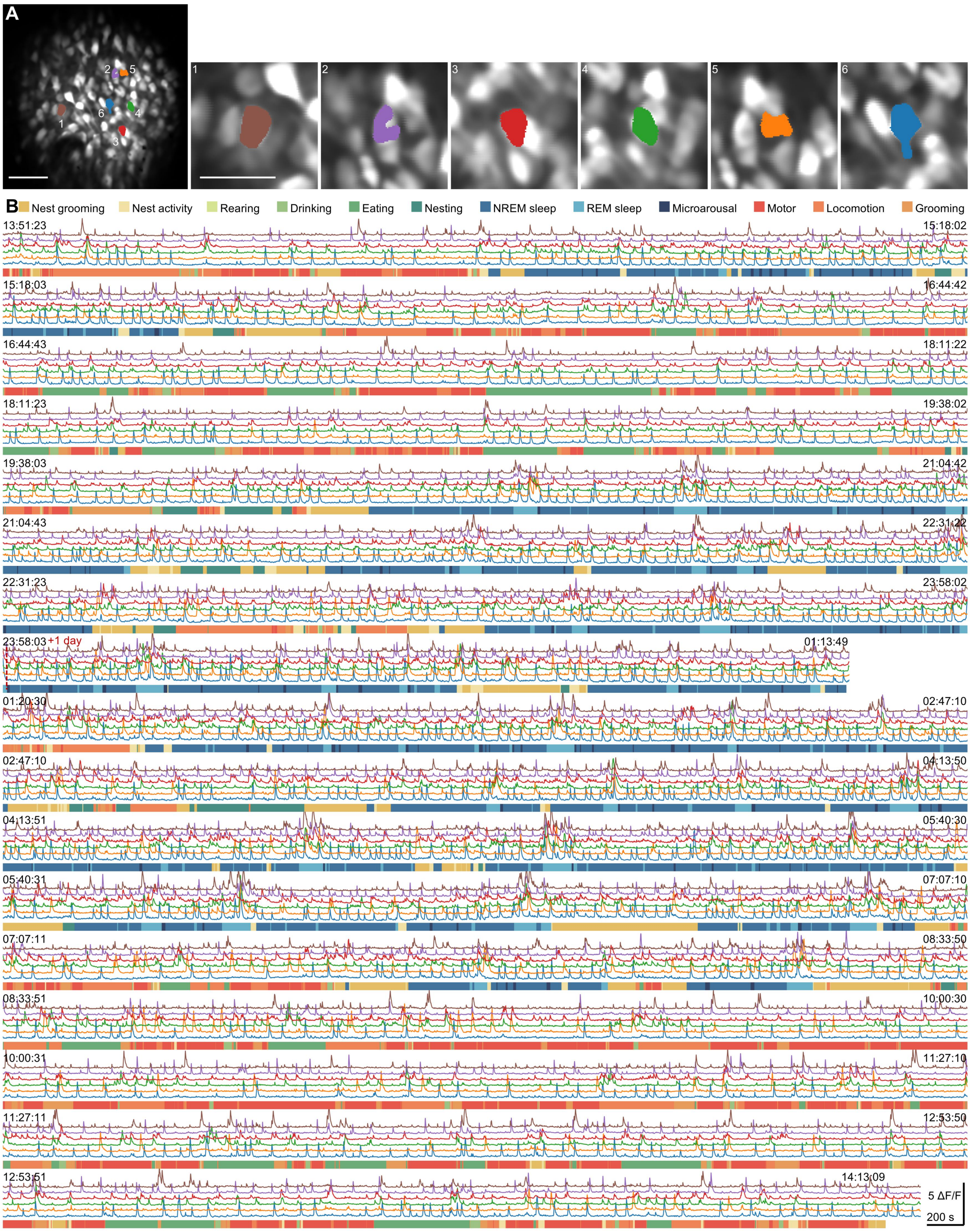


136 **Figure S6**

137 **24-hour mPFC neuronal activity traces aligned with spontaneous behavioral states**
138 **after NeuroPilot processing.**

139 (A) Representative FOV from 24-hour head-mounted two-photon calcium imaging in the
140 mPFC. Insets show magnified views of six selected neurons and their corresponding
141 segmented ROIs. Scale bars, 50 μm.

(B) Calcium activity traces from the six neurons shown in (A) across a 24-hour recording. Trace colors correspond to the ROI colors in (A). Successive rows show consecutive recording windows, with timestamps indicated above each window. Behavioral states are shown as a color-coded ethogram below each trace window. Scale bars, 5 $\Delta F/F$ and 200 s.

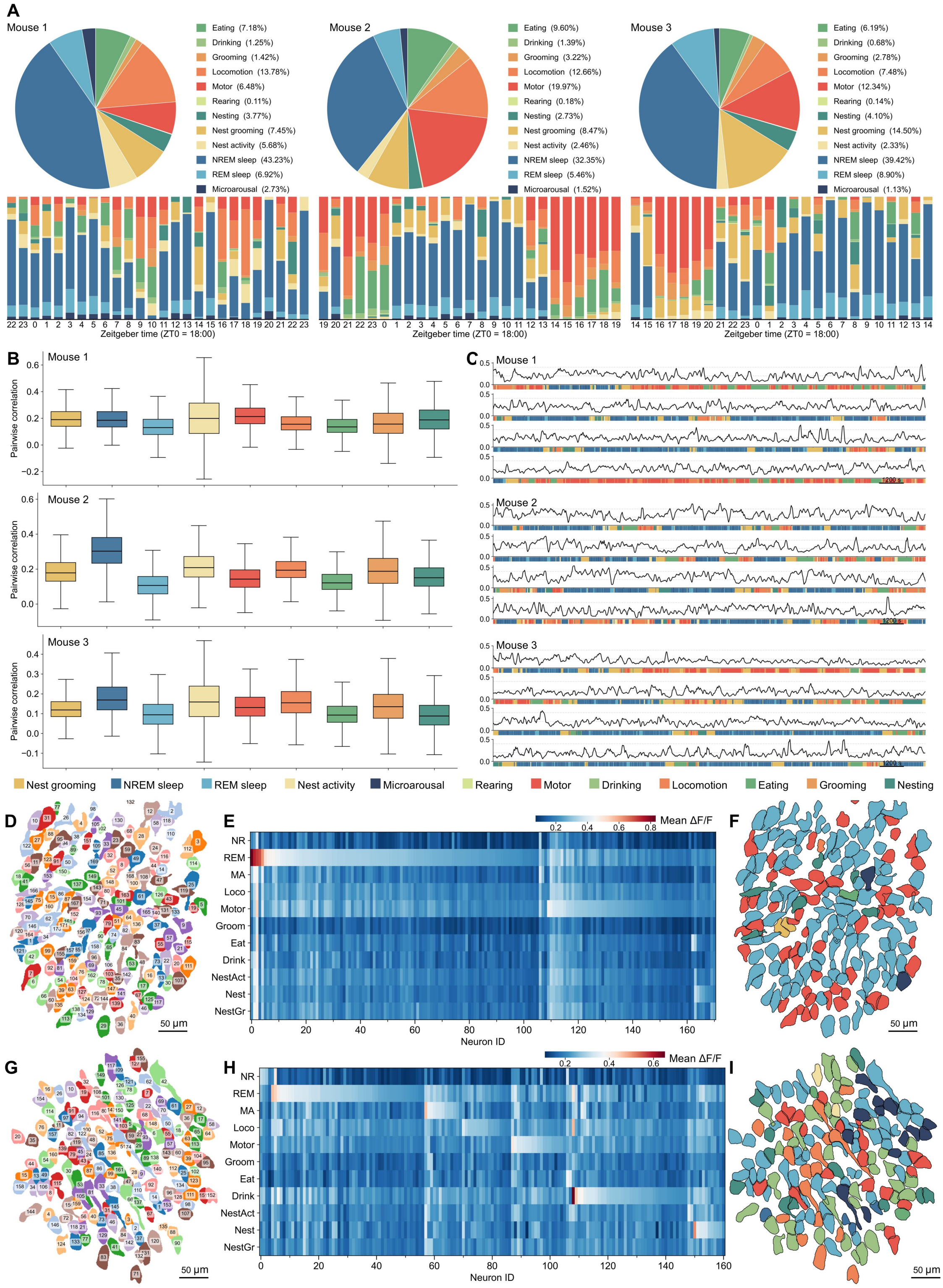


**Figure S7**

**Behavioral-state-dependent mPFC activity patterns and population correlations across 24-hour recordings.**

(A) Behavioral-state composition and Zeitgeber-time distribution for three representative mice. Top, pie charts show the fraction of recording time assigned to each behavioral state. Bottom, stacked bar plots show behavioral-state distributions across Zeitgeber time (ZT; ZT0 = 18:00). Each bar represents a 1 h bin, and colors indicate behavioral states.

(B) Pairwise population correlations grouped by behavioral state for mice 1, 2, and 3. Pearson correlation coefficients are computed from neuronal activity traces using 60 s sliding windows and grouped according to the behavioral state assigned to each window. Box plots summarize the resulting correlation distributions.

(C) Population-averaged pairwise correlation dynamics for all mice. Pairwise correlations are computed using 60 s sliding windows and averaged across neuron pairs. Successive traces show consecutive recording epochs, with aligned behavioral-state annotations below. Scale bars, 1,200 s.

(D-F) Behavioral-state-dependent activity patterns from mouse 1.

(D) Spatial map of segmented neuronal somata.

(E) Mean ΔF/F of individual neurons across behavioral states, ordered by the state with the highest mean ΔF/F.

(F) Spatial distribution of neurons color-coded by the state with the highest mean ΔF/F. n = 171 neurons.

(G-I) Same analyses as in (D-F) for mouse 3. n = 161 neurons.

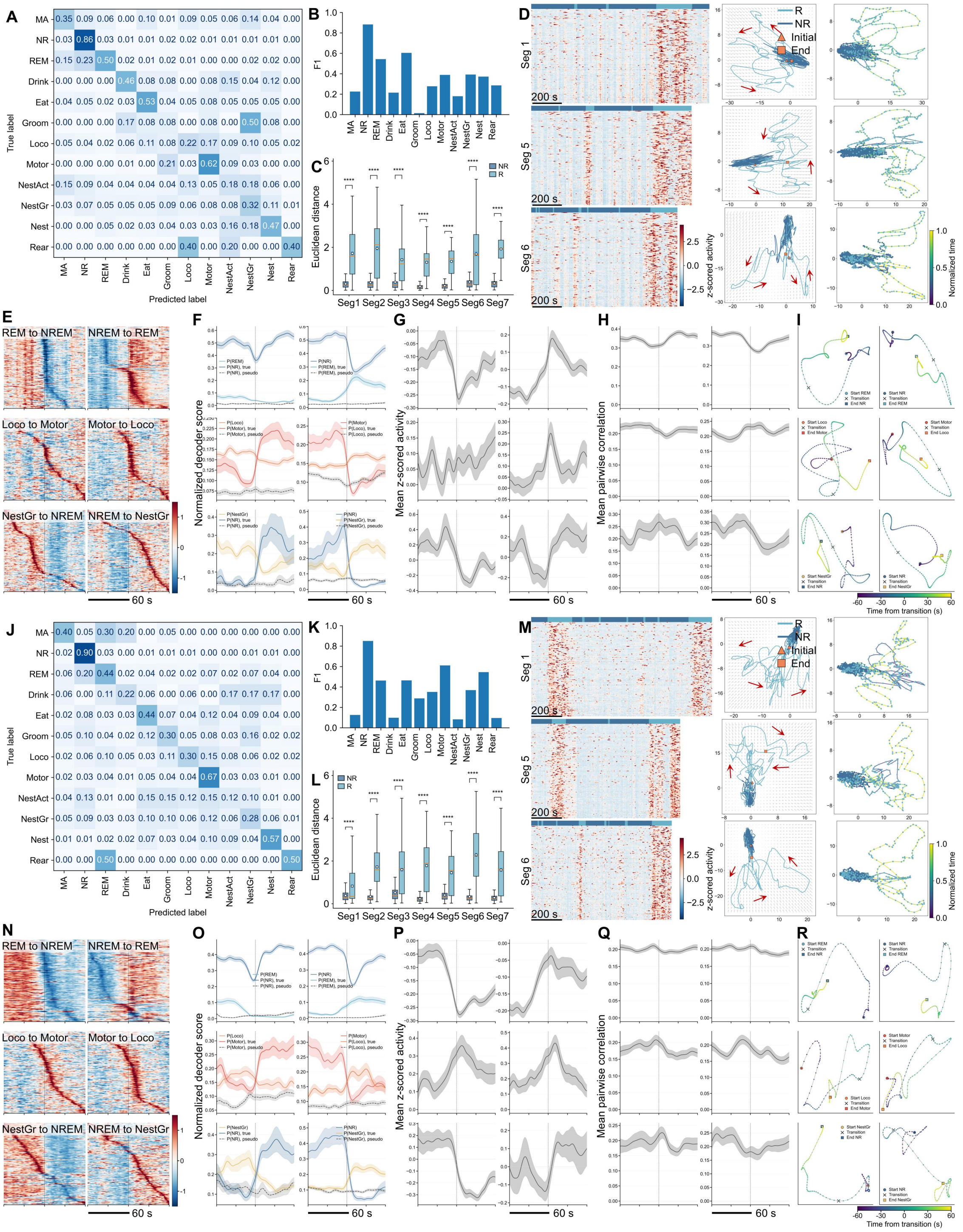


**Figure S8**

**Behavioral-state decoding and transition dynamics of mPFC population activity in additional mice.**

(A–I) Analysis of 24-hour mPFC population activity from mouse 1.

(A) Confusion matrix for decoding 12 behavioral states from mPFC population activity. Values indicate the fraction of samples from each true behavioral state assigned to each predicted state. MA, microarousal; NR, NREM sleep; REM, REM sleep; Drink, drinking; Eat, eating; Groom, grooming; Loco, locomotion; Motor, motor activity; NestAct, nest activity; NestGr, nest grooming; Nest, nesting; Rear, rearing.

(B) F1 scores for the decoded behavioral states shown in (A).

(C) Euclidean distance to the NREM centroid across representative REM/NREM segments. Box plots compare NREM and REM time points within each segment.

(D) Representative REM/NREM segments showing z-scored neuronal activity heatmaps, latent trajectories with inferred vector fields, and corresponding trajectories colored by normalized time. Triangles indicate initial points, squares indicate endpoints, and red arrows indicate trajectory direction. Scale bars, 200 s.

(E) Population activity heatmaps aligned to six representative behavioral transitions. Activity is z-scored. Scale bar, 60 s.

(F) Transition-aligned behavioral decoder scores for the same transitions shown in (E).

(G) Mean z-scored population activity aligned to the same transitions.

(H) Mean pairwise population correlation aligned to the same transitions.

(I) Low-dimensional population trajectories around representative behavioral transitions.

(J–R) Same analyses as in (A)–(I) for mouse 3.

Box plots show the median, interquartile range, and whiskers indicating the data range. Statistical significance is indicated as ****$P < 0.0001$.
